# Phosphate dysregulation via the XPR1:KIDINS220 protein complex is a therapeutic vulnerability in ovarian cancer

**DOI:** 10.1101/2020.10.16.339374

**Authors:** Daniel P Bondeson, Brenton R Paolella, Adhana Asfaw, Michael Rothberg, Thomas Skipper, Carly Langan, Alfredo Gonzalez, Lauren E Surface, Kentaro Ito, Mariya Kazachkova, William N Colgan, Allison Warren, Josh Dempster, Mike Burger, Maria Ericsson, Andrew Tang, Iris Fung, Emily S Chambers, Mai Abdusamad, Nancy Dumont, John G Doench, Federica Piccioni, David E Root, Jesse Boehm, William C Hahn, Michael Mannstadt, James M McFarland, Francisca Vazquez, Todd R Golub

## Abstract

Clinical outcomes for patients with ovarian and uterine cancers have not improved greatly in the past twenty years. To identify ovarian and uterine cancer vulnerabilities, we analyzed genome-scale CRISPR/ Cas9 loss-of-function screens across 739 human cancer cell lines. We found that many ovarian cancer cell lines overexpress the phosphate importer SLC34A2, which renders them sensitive to loss of the phosphate exporter XPR1. We extensively validated the XPR1 vulnerability in cancer cell lines and found that the XPR1 dependency was retained in vivo. Overexpression of SLC34A2 is frequently observed in tumor samples and is regulated by PAX8 -a transcription factor required for ovarian cancer survival. XPR1 overexpression and copy number amplifications are also frequently observed. Mechanistically, SLC34A2 overexpression and impaired phosphate efflux leads to the accumulation of intracellular phosphate and cell death. We further show that proper localization and phosphate efflux by XPR1 requires a novel binding partner, KIDINS220. Loss of either XPR1 or KIDINS220 results in acidic vacuolar structures which precede cell death. These data point to the XPR1:KIDINS220 complex - and phosphate dysregulation more broadly -as a therapeutic vulnerability in ovarian cancer.

---

Precision cancer medicine aims to develop therapies which can be given to individuals with tumors most likely to benefit from such a therapy. Although the discovery of these “ predictable dependencies” has significantly benefited many other tumor types, ovarian and uterine cancers have not seen as much benefit and the outcomes for these cancers has not significantly changed in the past twenty years^1,2^. To identify novel therapeutic targets for ovarian and uterine cancers, we analyzed genome-scale, pooled CRISPR/Cas9 loss of viability screens in 739 genomically characterized human cancer cell lines as part of the Cancer Dependency Map^3–5^. We focused on genes which, when inactivated, selectively led to loss of viability in ovarian or uterine cancer cell lines, since a broad killing pattern is more likely to represent mechanisms that would be poorly tolerated if pharmacologically inhibited. This analysis (Figure 1a) yielded ‘selective dependencies’ such as the transcription factor PAX86 and other genes with highly selective killing patterns but without a clear path toward therapeutic development. This analysis also revealed that inactivation of the phosphate exporter XPR1 has a cell-killing pattern which was highly selective and enriched in ovarian and uterine cancers. XPR1 is a viral receptor transmembrane protein7 and the only phosphate exporter annotated in human biology^8,9^. Direct inhibition of the phosphate efflux capacity of XPR1 is likely possible9, but its relevance in ovarian cancer has not been explored before.

**Figure 1:**
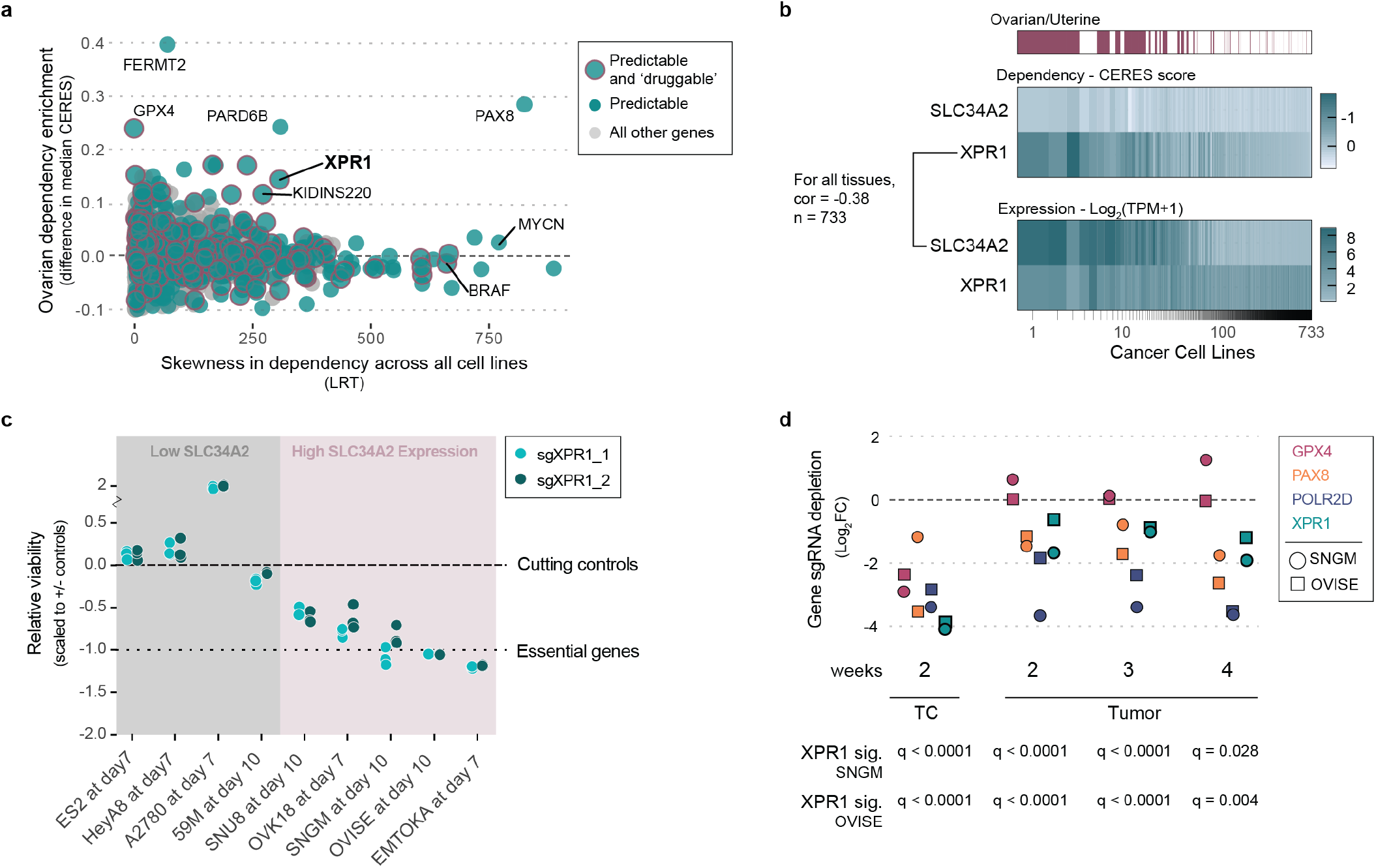
An ovarian cancer vulnerability to loss of XPR1 is predicted by SLC34A2 expresion. **a**. For the 18,000 genes tested in CRISPR/Cas9 loss of viability screens, the selectivity of the killing profile across all 739 cell lines (X-axis) and the enrichment of that gene’s dependency (CERES score) in ovarian and uterine cancers (Y-Axis) is plotted. Whether a gene’s dependency profile is predictable from molecular features (predictive strength > 0.4) is indicated by teal dots, and gene products predicted to be ‘druggable’ by traditional inhibitors are indicated with black outlines. **b**. Heatmap indicating XPR1 and SLC34A2 expression (Log2(TPM+1)) and dependency (CERES) values across all cell lines, approximately ranked by decreasing dependency on XPR1. The pearson correlation coefficient across all 733 cell lines is indicated. **c**. Viability effects after inactivation of XPR1, scaled such that a value of 0 represents the small viability effect of CRISPR/Cas9 genome editing and -1 represents loss of an essential gene. “ High SLC34A2 expression” indicates mRNA expression greater than 3 TPM. **d**. The median depletion of sgRNA targeting XPR1 or other genes is plotted for CRISPR/Cas9 competition tumor formation assays performed in ovarian (OVISE, squares) or uterine (SNGM, circles) cell lines. GPX4 is a metabolic dependency gene. PAX8 is a benchmark dependency in many ovarian cancers. POLR2D is a pan-essential positive control. Below, the significance of XPR1 depletion compared to the depletion of control sgRNA was calculated via t-test.

We next pursued the molecular basis of the selective dependency on XPR1. Using more than 100,000 molecular features of the cancer cell lines^10^, we built multivariate models -potential “ biomarkers” of response -to predict XPR1 dependency^11,12^. Remarkably, the feature which most robustly predicted XPR1 dependency was expression of the phosphate importer SLC34A2 (Figure 1b, pearson coefficient = -0.38 in all cell lines, -0.58 in ovarian and uterine cancer cell lines). SLC34A2 overexpression in ovarian cancer is well documented^13,14^ and was highly correlated with XPR1 dependency in cell lines from the ovarian clear cell, high grade serous, and endometrial adenocarcinoma lineages (Supplemental Figure 1b).

We validated the pooled screening results in a panel of SLC34A2 high-and low-expressing ovarian and uterine cancer cell lines, confirming that XPR1 is a selective dependency in the context of SLC34A2 overexpression (Figure 1c and Supplemental Figure 1c-e). We next assessed the XPR1 dependency in a CRISPR/Cas9-based tumor formation competition assay, and observed XPR1 sgRNA depletion in two SLC34A2-high tumors (Figure 1d and Supplemental Figure 2). In contrast, sgRNA targeting other metabolic dependencies, such as the ferroptosis regulator GPX4, were depleted in vitro but not in vivo, as has been previously reported (Figure 1d)^15,16^. Consistent with these results, preliminary data in a model of established peritoneal carcinomatosis suggested that suppression of XPR1 expression inhibits tumor progression (Supplemental Figure 3). Taken together, these results indicate that the XPR1 dependency is retained in vivo.

**Figure 2:**
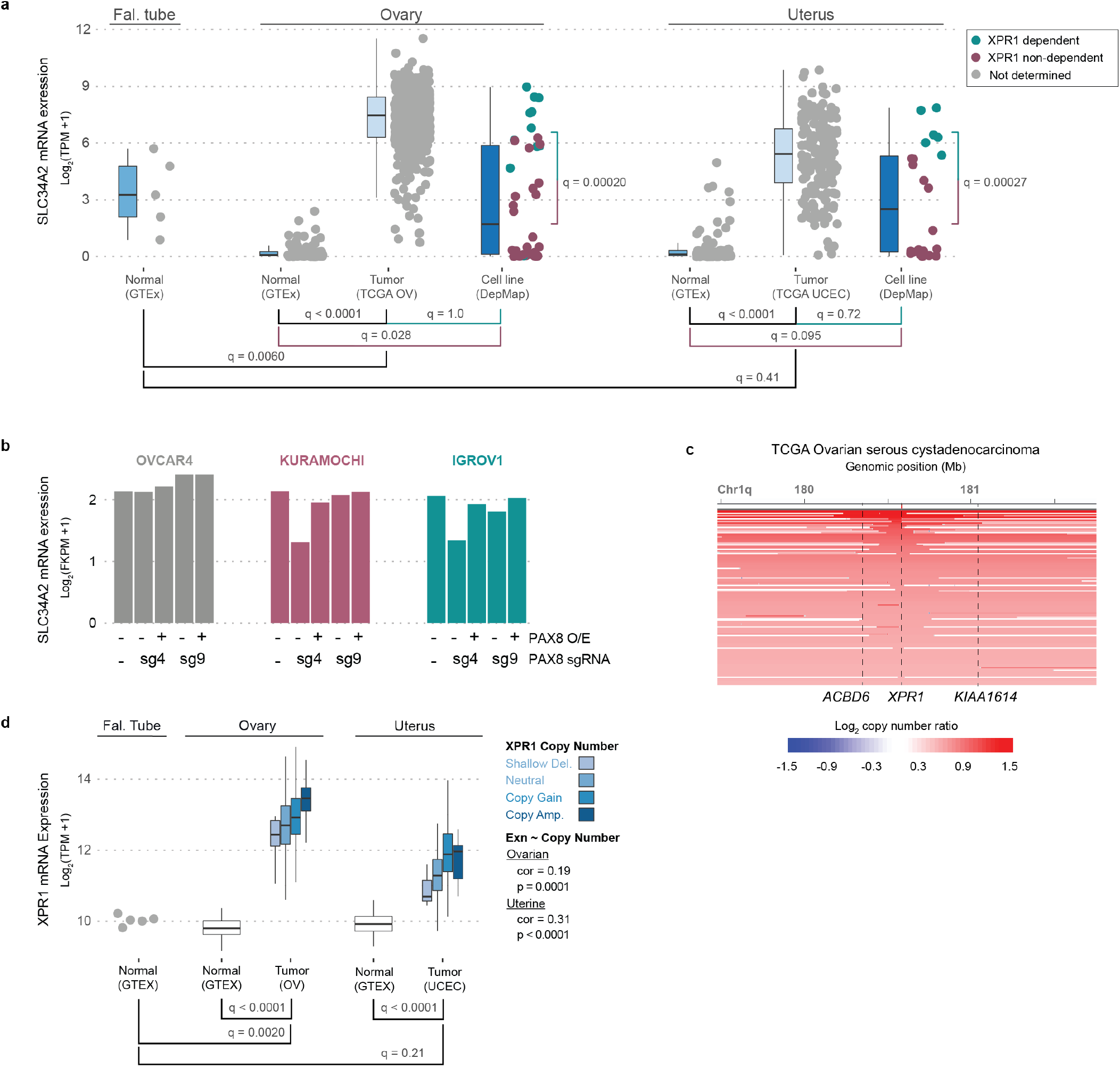
XPR1 and SLC34A2 expression in patient samples indicate cancer-specific phosphate dysregulation and a therapeutic opportunity for XPR1 inhibition. **a**. SLC34A2 mRNA expression is compared in normal (GTEx), tumor (TCGA), and cancer cell line (DepMap) samples. For CCLE cell lines, the XPR1-dependency status (CERES < -0.5) is indicated. Note that fallopian tube epithelium is the tissue of origin for many ovarian and uterine cancers. **b**. The expression of SLC34A2 was measured using RNAseq after stable overexpression of PAX8 as indicated, and induction of a PAX8-targeting (sg4) or control (sg9) sgRNA and dCas9-KRAB. **c**. XPR1 copy number heatmap for a ∼2.5 Mb region of chromosome 1 indicating XPR1 amplification in serous ovarian cancer (TCGA OV). **d**. As in panel a, XPR1 mRNA expression is compared across the indicated tissues. For TCGA samples, the XPR1 copy number status (GISTIC) is indicated and the spearman correlation between copy number status and mRNA expression is indicated on the right.

**Figure 3:**
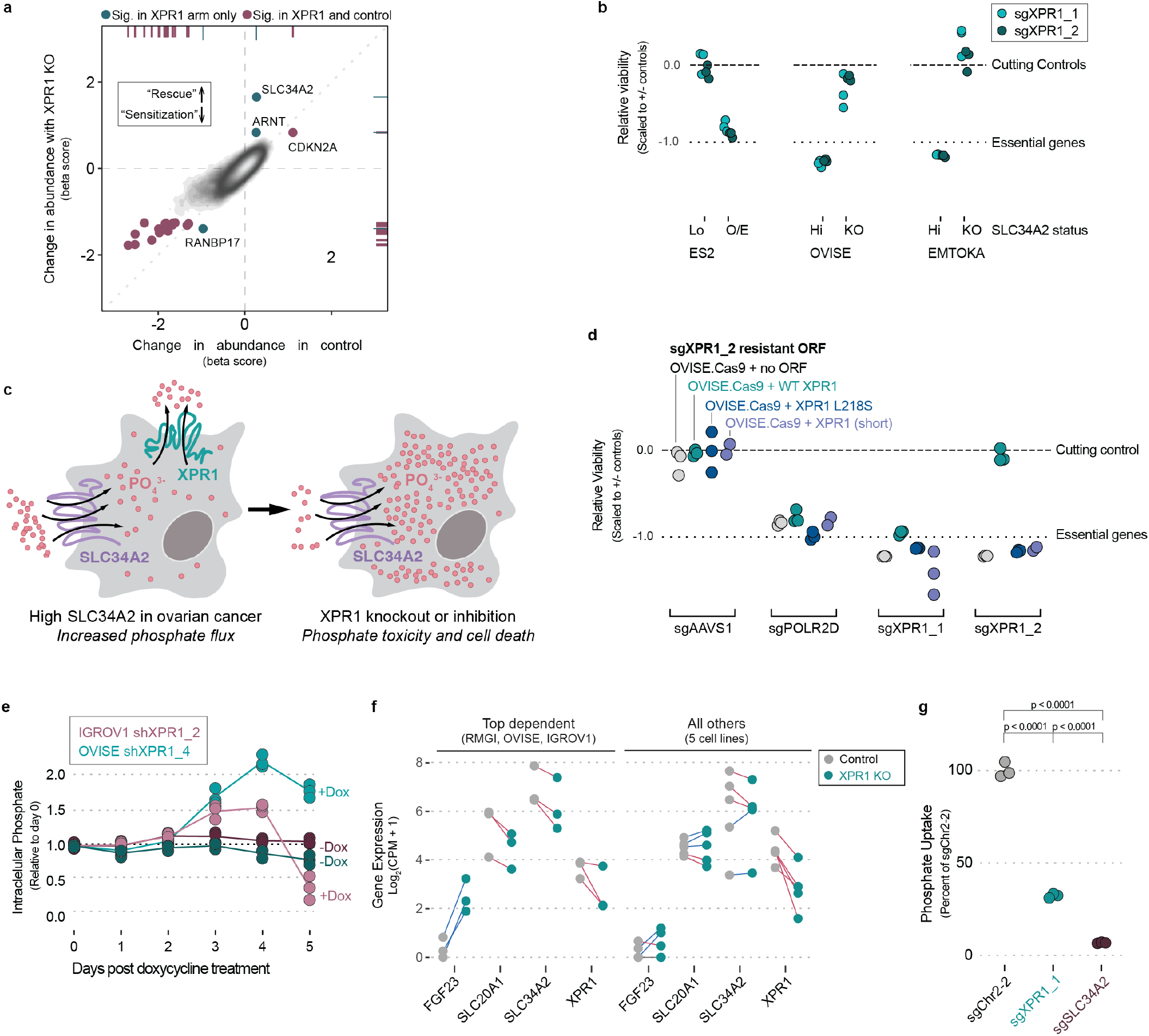
XPR1 inactivation in SLC34A2-high ovarian cancer causes cell death via dysregulated intracellular phosphate homeostasis. **a**. Genome-scale CRISPR/Cas9 screen combined with inactivation of XPR1 to find potential ‘modifier’ genes for the dependency. Beta scores (determined by MaGeCK MLE) represent the change in representation for each gene from the initial library to the final timepoint for the control condition (X-axis) or combined with XPR1 inactivation (Y-axis). **b**. The SLC34A2 status of normally XPR1-resistant (ES2, SLC34A2-low [lo]) or XPR1-sensitive (EMTOKA and OVISE, SLC34A2-high [hi]) cell lines was modified by overexpression (O/E) or inactivation (KO) of SLC34A2, and the XPR1 dependency was evaluated as in Figure 1c. **c**. Because of their relative directionalities of phosphate transport, we hypothesize that XPR1 inactivation is toxic because of intracellular phosphate accumulation in SLC34A2-high ovarian cancer. **d**. The indicated XPR1 open reading frames were tested for their ability to rescue inactivation of endogenous XPR1. The L218S mutation in XPR1 has previously been shown to only have ∼50% the phosphate efflux function of wild-type XPR1 (see main text). **e**. At various time-points after treatment with doxycycline and induction of shRNA, the intracellular phosphate was measured in OVISE and IGROV1 cell lines. **f**. The measured transcriptional change in the indicated genes are plotted for the cell lines with the largest and most correlated transcriptional change (see Supplemental Figure 7) on the left and the other five cell lines on the right. Red lines connect cell lines displaying increased expression upon XPR1 inactivation, blue lines indicate decreased expression. **g**. Radioactive phosphate uptake was measured in the OVISE ovarian cancer cell line after inactivation of the indicated genes.

To extend the relevance of XPR1 and SLC34A2 beyond cell lines, we evaluated the relationship between XPR1 and SLC34A2 in primary patient samples from The Cancer Genome Atlas (TCGA)^17,18^ and normal samples from the Genotype-Tissue Expression project (GTEx)^19– 21^. Ovarian tumors, on average, expressed SLC34A2 at levels 16-fold higher than normal fallopian tube epithelium (q = 0.006), which is thought to be the cell of origin of ovarian cancers^22–25^ (Figure 2a). Uterine cancers similarly overexpressed SLC34A2 relative to normal tissue. Interestingly, ovarian and uterine cancers were among the few tissues with high levels of SLC34A2 expression (Figure 2a and Supplemental Figure 4a). We hypothesized that SLC34A2 expression may be maintained at high levels in ovarian cancer because of its regulation by the transcription factor PAX8, the expression of which is elevated in ovarian cancer and required for its survival^6,26^. In support of this, we found a strong correlation of PAX8 and SLC34A2 expression in patient samples (Supplemental Figure 4b,c), and CRISPR interference-mediated downregulation of PAX8 led to a significant loss of SLC34A2 expression and other previously reported PAX8 target genes^27^ (Figure 2b and Supplemental Figure 4d).

**Figure 4:**
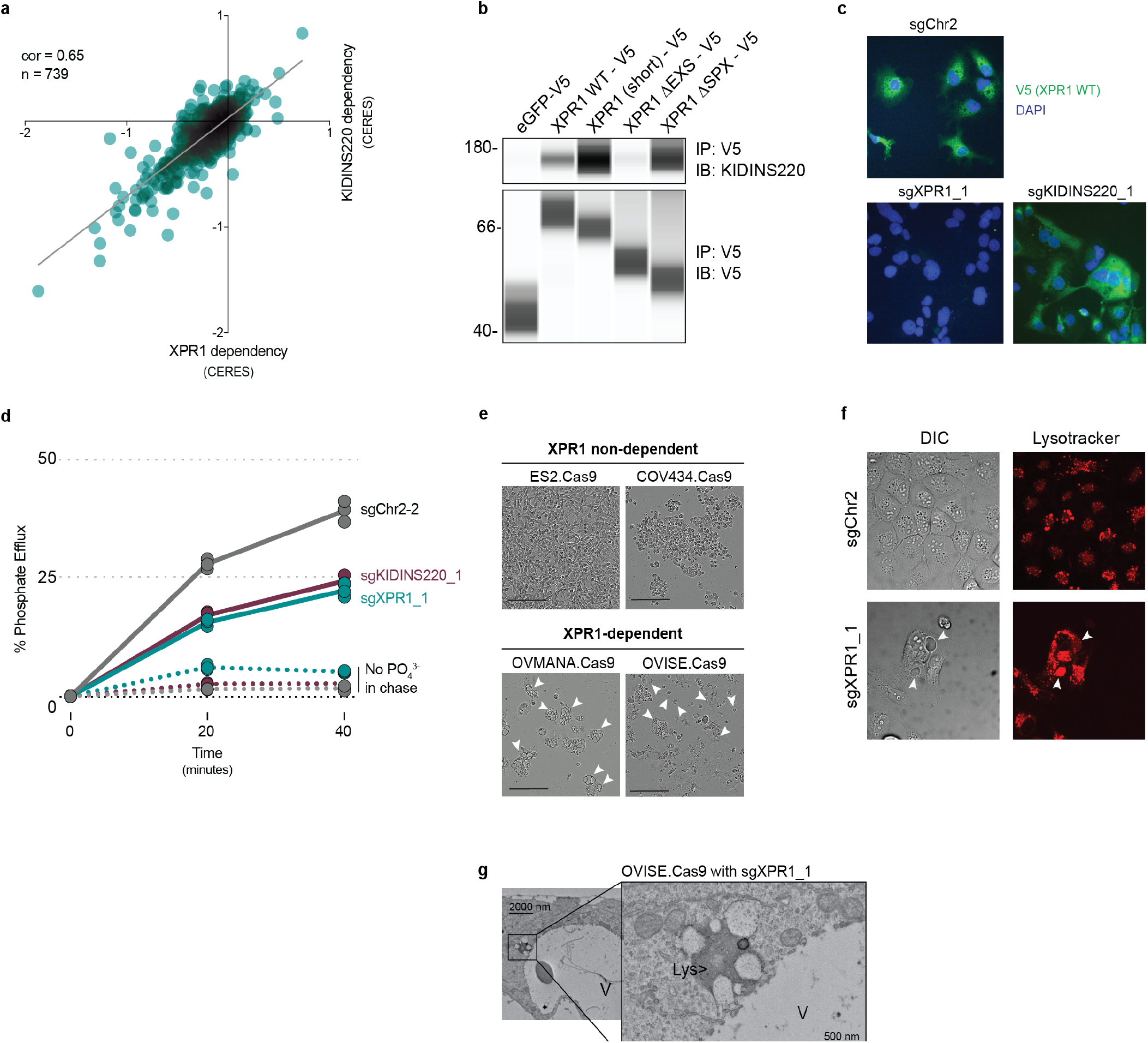
Protein efflux is achieved by the XPR1:KIDINS220 protein complex and perturbation of this complex causes accumulation of acidic ‘vacuole’ structures. **a**. Across 739 cancer cell screened (DepMap Avana 20Q1), the viability defects of XPR1 or KIDINS220 inactivation is plotted and the pearson correlation is indicated. **b**. After expression of the indicated XPR1 open reading frames in HEK-293Ts, the interaction between the V5-tagged ORF and KIDINS220 was evaluated using co-immunoprecipitation. See Supplemental Figure 9a for the design of these constructs. **c**. After inactivation of XPR1 or KIDINS220, the localization of XPR1-V5 was determined using immunofluorescence. Note that sgXPR1_1 inactivates both endogenous XPR1 and the XPR1-V5 ORF, and so no staining should be expected. **d**. Three days after genetic inactivation of XPR1 or KIDINS220, cells were loaded with radioactive phosphate, and then efflux was measured for 30 minutes after. **e**. Phase-contrast images of ‘vacuole-like’ phenotype 4-5 days after XPR1 inactivation. Arrowheads indicate the location of ‘vacuole-like’ structures. Scale bars = 200 μm. **f**. The acidic dye Lysotracker was used to stain live cells five days after inactivation of XPR1. **g**. Transmission electron micro-graphs of “ vacuole-like” structures (labeled V) or lysosomes (Lys) in OVISE cancer cells after *XPR1* inactivation.

In contrast to the more restricted tissue expression of SLC34A2 and PAX8, XPR1 is widely expressed in normal and cancer tissues (Supplemental Figure 4e). Nevertheless, we found strong evidence for positive selection of XPR1 copy number amplifications and enhanced mRNA expression in ovarian and uterine cancer, consistent with its dependency in these tissues (Figure 2c). In ovarian cancer, these amplifications were often focal, involving only the XPR1 gene (Figure 2d, q = 0.0015^28^) whereas in uterine cancer, broader and less significant amplifications were observed (Supplemental Figure 4f, q = 0.568). XPR1 mRNA expression levels were correlated with XPR1 copy number alterations, but other mechanisms likely also contribute to XPR1 mRNA expression (Figures 2d). Thus, the dysregulated expression of the essential transcription factor PAX8 results in enhanced expression of SLC34A2, thereby creating dependency on XPR1.

To further understand the mechanism by which XPR1 loss-of-function results in cancer cell death, we performed a genome-wide CRISPR/Cas9 rescue screen to determine which genes, when inactivated, were capable of rescuing XPR1-mediated loss of viability (Figure 3a, Supplemental Figure 5)^29^. Remarkably, the top rescuing gene (>18,000 tested), was also the top predictive biomarker: SLC34A2. We further demonstrated that SLC34A2 is both necessary and sufficient to confer XPR1 dependency in ovarian and uterine cell lines (Figure 3b).

The observation that high expression of the phosphate importer SLC34A2 is required for cell death after inactivation of the phosphate exporter XPR1 led us to hypothesize that accumulation of intracellular phosphate is toxic to ovarian and uterine cancer cells (Figure 3c). XPR1 is the only known phosphate exporter in humans^9^, suggesting that in the context of increased phosphate import, efflux would be in higher demand and require XPR1 function. Although the extracellular availability of phosphate in rich tissue culture medium far exceeds what is physiologically relevant, we found no correlation between the phosphate content of growth medium and XPR1 dependency across the Cancer Dependency Map dataset (Supplemental Figure 6a). Furthermore, XPR1 dependency was retained when cells were adapted to growth medium with near-physiological phosphate concentrations (reduced by ∼90% from 72.8 to 7.8 mg/dL, Supplemental Figure 6b-c), indicating that XPR1 dependency is not an artifact of the high concentrations of extracellular phosphate common in tissue culture growth medium. We also confirmed that the phosphate efflux function of XPR1 was critical for cell survival. Expression of a naturally occuring hypomorphic XPR1 mutation (L218S), associated with a rare brain calcification disorder^30^, failed to rescue endogenous XPR1 inactivation, whereas wild-type XPR1 fully restored cell viability (Figure 3d and Supplemental Figure 6d-e).

Consistent with the phosphate accumulation hypothesis, we observed 2-4 fold increased intracellular phosphate after XPR1 suppression (Figure 3e). These large fluctuations in intracellular phosphate co-occur with loss of cell viability (Supplemental Figure 7a)^31^. To understand the cellular response to phosphate accumulation, we used single-cell RNA sequencing of 2,501 cells across 8 ovarian and uterine cancer cell lines at an early time-point following XPR1 inactivation (Supplemental Figure 7b-i)^32^. The resulting transcriptional signature indicated cellular attempts to restore phosphate homeostasis, including the up-regulation of FGF23 (Figure 3f). This critical phosphate homeostatic hormone is typically expressed in osteogenic bone cells, and its expression in ovarian cancer cells -although not represented at the protein level (Supplemental Figure 7j) -is consistent with sensing elevated phosphate^33^. We also observed downregulation of phosphate importers at both the mRNA (Figure 3f) and protein level (Supplemental Figure 1b) after XPR1 inactivation. Consistent with this, XPR1 inactivation led to a 60% decrease in phosphate uptake (Figure 3g), suggesting that phosphate uptake is regulated by intracellular phosphate concentrations^34,35^. We note, however, that this partial compensatory response is not sufficient to protect cells from XPR1-mediated cell death.

To gain further insight into the mechanism by which XPR1 regulates phosphate homeostasis, we analyzed the Cancer Dependency Map for genes with highly correlated dependency profiles to XPR1. These “ co-dependencies” often indicate proteins that are part of the same protein complex^36,37^. Of the ∼18,000 genes analyzed, XPR1 dependency is most strongly correlated with KIDINS220, a gene with no known connection to phosphate homeostasis^38–41^ (pearson correlation = 0.81, Figure 4a and Supplemental Figure 8a). Given the strength of this correlation, we extensively validated KIDINS220 dependency (Supplemental Figure 8b-c), and hypothesized that KIDINS220 might be part of an XPR1 phosphate export complex.

In support of an XPR1:KIDINS220 protein complex, protein interaction databases indicate XPR1 and KIDINS220 interact with each other (Supplemental Figure 9a). Further, their gene expression was highly correlated across diverse tissues (Supplemental Figure 9b), suggesting co-function and co-regulation. To confirm this interaction, we performed co-immunoprecipitation experiments and found that XPR1 and KIDINS220 indeed form a protein complex (Figure 4b, Supplemental Figure 9c-d). We mapped the XPR1:KIDINS220 interaction to the C-terminus of XPR1 containing the EXS domain, an evolutionarily conserved domain known to be required for XPR1 trafficking between the golgi apparatus and the plasma membrane in order to achieve phosphate efflux^42–44^. In contrast, the N-terminal SPX domain of XPR1, which has been implicated in phosphate efflux and regulation^35,45^, was neither necessary nor sufficient to bind KIDINS220 (Supplemental Figure 9b).

Further supporting an XPR1:KIDINS220 protein complex, we found dramatically decreased KIDINS220 protein levels following XPR1 suppression (Supplemental Figure 1b). In addition, KIDINS220 inactivation caused XPR1 to mislocalize from punctate secretory vesicles to a more diffuse pattern (Figure 4c). Finally, we directly measured phosphate efflux and found that inactivation of either XPR1 or KIDINS220 impaired phosphate efflux (Figure 4d) and resulted in increased intracellular phosphate (Supplemental Figure 9e). These results, taken together, indicate that phosphate efflux is achieved by the XPR1:KIDINS220 protein complex, and that loss of either complex member leads to a disruption in phosphate efflux which is required for cancer cell survival.

A striking feature of XPR1- or KIDINS220-mediated cell death is the formation of large cytoplasmic structures resembling vacuoles immediately preceding loss of cell viability (Figure 4e-g, Supplemental Figure 10a). While a panel of organellar stains did not identify these structures (Supplemental Figure 10b), co-localization with the acidic dye Lysotracker and the lysosomal marker LAMP1 (Figure 4f and Supplemental Figure 10b) suggested they may be related to the lysosomal system. Ultrastructural analysis by transmission electron microscopy found these structures to be bound by a double-membrane that was often fenestrated (Figure 4g and Supplemental Figure 10d). Although they lack the electron-dense appearance typical of lysosomes, we did note their fusion with lysosomes.

In yeast and plants, excessive uptake and defective efflux of phosphate leads to its storage as polyphosphate -long chains of inorganic phosphate -in vacuolar structures. Storage mechanisms for inorganic phosphate in human cells are yet to be discovered, but there are emerging studies indicating that the metabolism of phosphate, ATP, inositol pyrophosphates, and polyphosphate are all interconnected^8, 46-48^. Whether the structures observed after XPR1 inactivation in SLC34A2-overexpressing ovarian cancer cells reflect a compensatory cell survival mechanism by sequestering toxic phospho-metabolites, or are themselves the cause of cell death remains to be determined.

The study reported here highlights a previously unappreciated strategy to kill cancer cells: the disruption of phosphate homeostatic mechanisms that are normally tightly regulated. In ovarian cancer in particular, dysregulated expression of SLC34A2 by the essential transcription factor PAX8 abrogates normal homeostatic mechanisms and creates a potential therapeutic window. In support of this, XPR1 dependency was rarely observed in tissues with normally high levels of SLC34A2 expression (e.g. lung, Supplemental Figure 1b). Thus, under conditions of high SLC34A2 expression, loss of XPR1 or KIDINS220 leads to the intracellular accumulation of inorganic phosphate, which has previously been shown to be toxic to cells^49,50^. The experiments described here thus provide a rationale for the development of therapeutic strategies aimed at inhibiting XPR19 or KIDINS220^51^ in ovarian cancer, and more generally highlight the potential of disrupting phosphate homeostasis as a therapy for cancer.

## Acknowledgments

This work was funded in part by the Slim Initiative in Genomic Medicine for the Americas (SIGMA), a joint U.S-Mexico project funded by the Carlos Slim Foundation (TRG), grants from the National Cancer Institute CA242457 (TRG) and CA212229 (DPB), and the Robertson Foundation (TRG). We thank J Barnett, B Buckley, and M Veneskey for technical support.

## Author Contributions

DPB, BP, FV, and TRG initiated the project and oversaw the research plan. MB, DPB, and BP analyzed genetic dependency data under the supervision of WCH, DER, JB, FV, and TRG and support from IF and EC. MR, AA, TS, BP, and DPB conducted viability experiments and immunoblotting. AA, BP, and DPB conducted the genome-scale modifier screen with analysis support of MK, JD, and JMM and supervision from JD. In vivo experiments were conducted by AG and ND under the supervision of FP. Intracellular phosphate assays were performed by MR and DPB. PAX8 RNAseq experiments were conducted by KI with analytical support from WC. DPB analyzed GTEx, TCGA, and CCLE expression datasets with supervision from JMM. Multiplexed transcriptional profiling was conducted by BP and DPB and was analyzed by AW and WC. Phosphate uptake and efflux assays were conducted by DPB and LES with supervision from MM. AA and DPB conducted co-immunoprecipitation experiments. BP conducted cellular imaging studies. DPB and ME conducted ultrastructural analysis. DPB, BP, FV, and TRG wrote the manuscript and all authors edited and approved the manuscript.

## Tables

Supplemental Table 1: Gene-level beta scores and significance for the genome-scale CRISPR/Cas9 loss of function screen in combination with XPR1 inactivation.

Supplemental Table 2:sgRNA sequences used in the in vivo competition assay (related to Figure 1g).

## Data Availability

Publicly available data used in this study include CRISPR/Cas9 loss of viability screens for 739 cancer cell lines^3–5^, cancer cell line RNAseq expression data^10^, harmonized gene expression data for GTEx and TCGA datasets^19^, and copy number alterations for Ovarian adenocarcinoma and Uterine Corpus Endometrial Carcinoma^17,18^. RNA sequencing data after PAX8 suppression and multiplexed transcriptional profiling are available from the corresponding author upon request.

## Code Availability

Computer code to reproduce these results is available from the corresponding authors upon request.

## Materials and Methods

### Genetic Dependency Data

The dependency data used in this manuscript come from the Public Avana 20Q1 dataset consisting of dependency data for 18,333 genes across 739 cancer cell lines from 26 lineages. Expression data from the Cancer Cell Line Encyclopedia was also used. These data are available online at depmap.org/downloads. In Figure 1a, the selectivity (NormLRT^52^) and predictability^5,12^ was determined as previously reported. “ Highly Predictable” genes are indicated if the pearson correlation coefficient between the experimental data and the top predictive model is greater than 0.4. The median dependency for each gene in ovarian/uterine cancers (n = 62) was subtracted from the median dependency in all other cancer cell lines (n = 671) to calculate the ovarian/ uterine genetic dependency enrichment on the Y-axis of Figure 1a. In Figure 1b and Supplemental Figure 1b, the correlation of SLC34A2 expression and XPR1 dependency was performed using a two-tailed pearson correlation test. To compare XPR1 co-dependencies (Supplemental Figure 8a), a two-tailed pearson correlation test was performed for XPR1 versus all other genes (k = 18,333 genes, n = 739 cell lines although some cell line:gene pairs are not represented), and p-values are reported after correcting for multiple comparisons with the Benjamini-Hochberg method.

### Cell Lines

ES2, HeyA8, A2780, 59M, SNU8, OVK18, SNGM, OVISE, EMTOKA, IGROV1, OVCAR4, KURAMOCHI, RMGI, COV413a, JHOS4, HEC6 and JHUEM1 cancer cell lines were collected by the CCLE before distribution for our use. The sources of the aforementioned cell lines can be found at DepMap.org. All cell lines were adapted to growth in RPMI 1640 (Corning) + 10% FBS before use.

### sgRNA sequences

The negative control guides sgChr2 and sgAAVS1 were designed to cleave a gene desert and an intronic region in PPP1R12C, respectively, to control for the effects of DNA double-strand breaks. sgLacZ targets a sequence not found in the human genome. Positive control sRNA target common-essential splicing factors (SF3B1), ribosomal subunits (POLR2D), or kinesin motor proteins (KIF11). The 20 bp targeting sequences are:

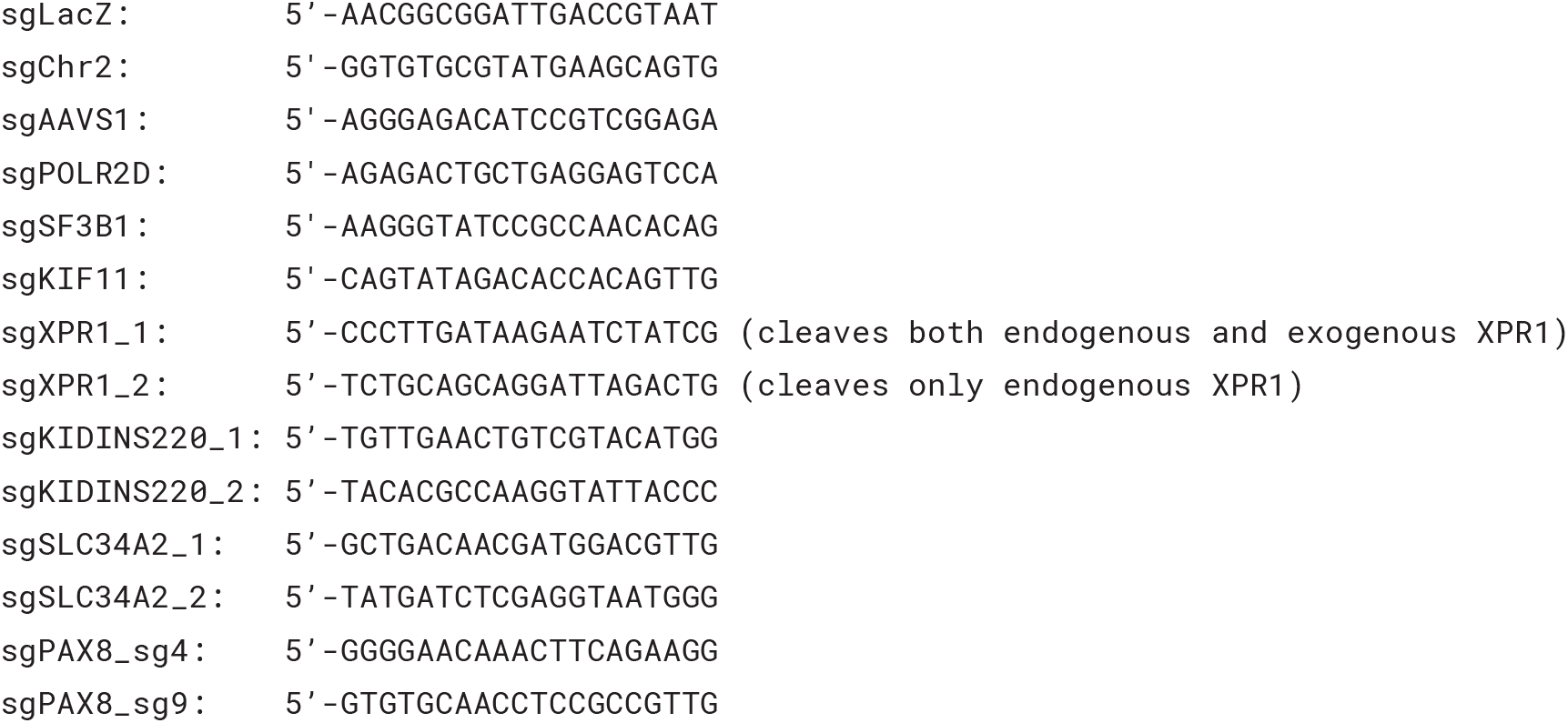

Additional guides included in the pooled *in vivo* knockout experiment are included in Supplementary Table 2.

### Lentiviral production

Lentiviral production was performed using HEK293T cells as described on the GPP portal (<u>GPP Web Portal -Home</u>).

### Plasmids, overexpression constructs, site directed mutagenesis

Open reading frames of the following genes were obtained from a genome-scale library of annotated genes^53^. SLC34A2 (NM_006424) was isolated from this library in pDONR223 and was transferred into the expression vector pLX-TRC313 (similar to Addgene #118017) using gateway cloning. The resultant construct has a C-terminal V5 tag, and after stable integration into cell lines using lentiviral infection, the proper protein product with a V5 tag was detected using wesern blot (not shown). XPR1 constructs (both isoforms NM_004736 and NM_001135669) were obtained in a similar way. Only the NM_004736 isoform was observed using isoform-agnostic PCR primers and cDNA generated from the OVISE cancer cell line. Mutations were introduced with PCR based methods, either the Q5 Site-directed Mutagenesis kit (NEB catalog E0554S) for large deletions or the QuickChange II XL (Agilent Catalog 200521) for point mutations, and were confirmed using sanger sequencing. The XPR1 ORFs were then transferred to pLX-TRC313 or the same expression vector with a weaker, PGK promoter. For co-immunoprecipitation experiments in HEK-293T cells, the stronger promoter XPR1 mutants were used to maximize expression levels. The weaker promoter construct was used for stable expression in ovarian cancer cell lines and immunofluorescence and mutant-rescue experiments.

### CRISPR Viability Assays

CRISPR viability assays were performed in 96-well plates with cells seeded at a low density to allow for logarithmic growth throughout the entire assay. For 7 day assays, cells were seeded and infected with lentivirus expressing the sgRNA in pXPR-BRD003 on Day 0. The next day, the infection media was replaced with 100 uL of media. On Day 7 post-infection, viability was evaluated by addition of 25 uL of Cell Titer Glo (Promega) reagent per well, and luminescence was measured. Infection efficiency was determined by comparing the viability of cells with and without puromycin after infection, and assays were repeated if less than 80% of cells were infected with every sgRNA. The data were normalized such that the cutting control sgRNA (targeting Chr2-2 and AAVS1) is 0 and positive control sgRNA (targeting the common essential genes KIF11, SF3B1, and POLR2D) is -1.0. For 10 day assays, infections were carried out in 6-well plates. Three days post-infection, the cells were lifted and seeded into replicate 96-well plates. On Days 3, 7 and 10 post-infection, viability was evaluated by addition of 50 uL of Cell Titer Glo (Promega) reagent per well and monitoring the luminescence. The fold change viability was calculated by comparing Days 7 or 10 to Day 3, and the data was normalized as above.

### shRNA sequences

shRNA sequences for XPR1 were selected from project DRIVE’s sub-genome scale shRNA library^52^ using DEMETER2 estimates for on-target and off-target seed effects 54. A detailed protocol for selecting shRNA using these datasets is available online (https://protocols.io/view/shrna-selection-and-quality-control-for-cancer-tar-bfmnjk5e). Doxycycline inducible shRNA expression was accomplished by cloning these sequences into the pRSITEP-U6Tet-(shRNA)-EF1-TetRep-2A-Puro vector (Cellecta #SVSHU6TEP-L). shRNA seed matched negative control sequences 55 were generated by substituting complementary base pair sequences into positions 9-11 bp of the target shRNA using the web-based tool (https://web.archive.org/web/20180605134130/http://rnai.nih.gov/haystack/C911Calc2.html).

shXPR1 and seed matched control sequences were rigorously tested for on-target XPR1 suppression as well as off target cell viability effects. Off target cell viability effects were determined when a given shSeed control sequence did not knockdown XPR1 but produced strong loss of cell viability, regardless of whether a cell line was predicted to be XPR1 dependent or non-dependent. shRNA target sequences are provided below:

~~~
shXPR1_2: 5’-CGCAGGTTTGCTACACTTCAG
shSeed_2: 5’-CGCAGGTTACGATCACTTCAG
shXPR1_4: 5’-CCTTGTGCTTGCCGCTGTATT
shSeed_4: 5’-CCTTGTGCAACCCGCTGTATT
~~~

### Antibodies

The following antibodies were purchased from the indicated sources. XPR1, KIDINS220, and SLC34A2 antibodies were validated by western blot of cell extracts after knockout or overexpression of the indicated gene. PS: Protein Simple. WB: Western Blot. IF: Immunofluorescence.

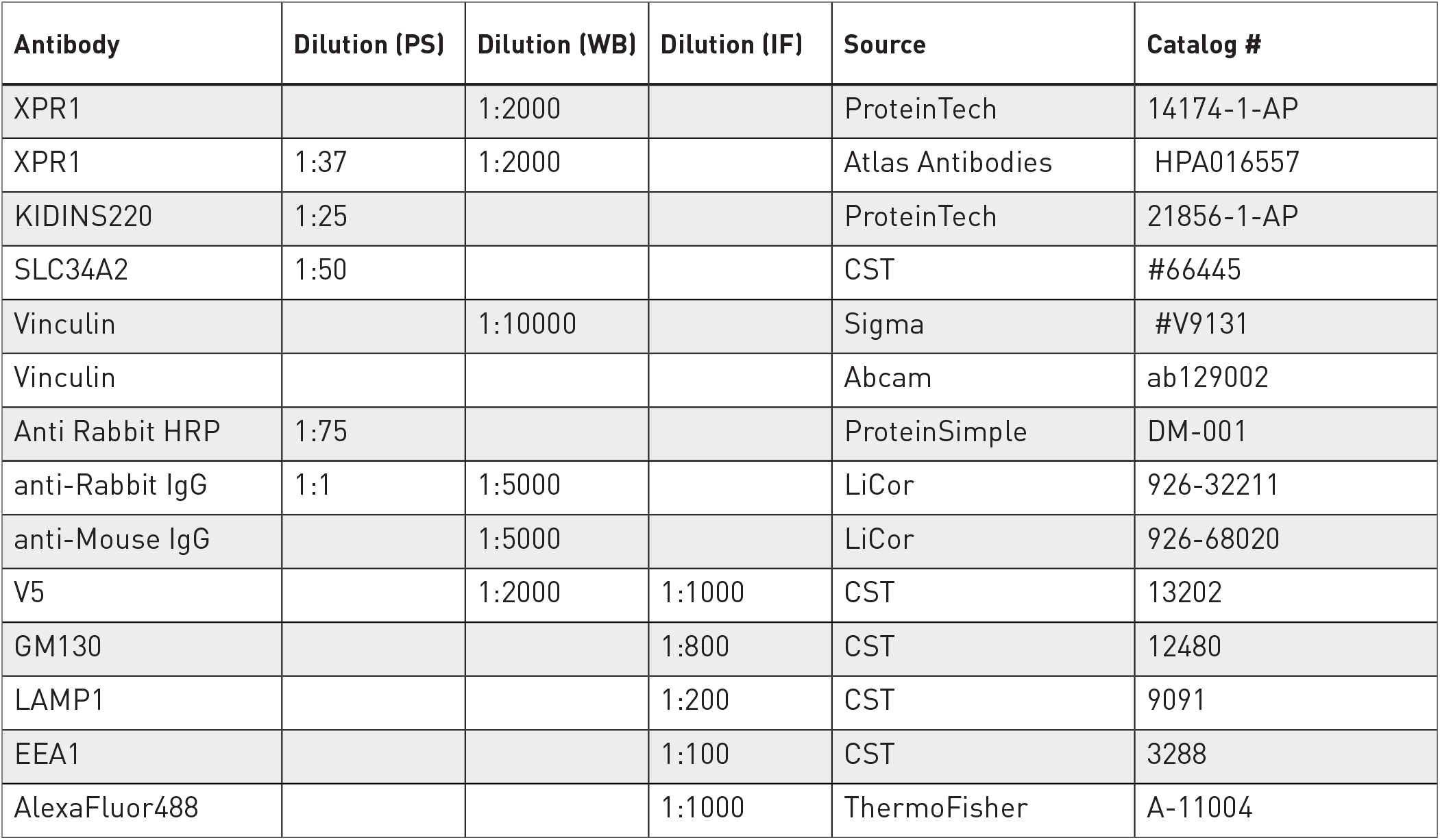

### Protein analysis of cell lysates by immunoblotting

For protein analyses, cells were grown in 6-well dishes. The cells were harvested by washing with PBS, incubating with 0.5 mL of TrypLE until all cells had lifted and diluting to 1.5 mL with PBS. The cells were then centrifuged and washed once with PBS, then lysed with Radioimmunoprecipitation assay (RIPA) buffer (150 mM NaCl, 1.0% IGEPAL® CA-630,0.5% sodium deoxycholate, 0.1% SDS, 50 mM Tris, pH 8.0) supplemented with cOmplete, Mini Protease and Phosphatase Inhibitor Cocktail Tablets (Roche). Cell extracts were cleared by spinning at 15000 rpm for 10 minutes at 4°C. Protein content was quantified by BCA analysis and 4x LDS reducing sample buffer was added. We found boiling the lysates led to a loss of XPR1 protein by western blot, and so the lysates were incubated at 37°C for 30 minutes to denature proteins. For standard western blots (i.e. Supplemental Figure 5b and Supplemental Figure 6e), equal amounts of protein (typically 25 μg) were resolved by SDS-PAGE, transferred to nitrocellulose and incubated with the indicated primary and secondary antibodies were used to visualize protein levels. Images were obtained using a Licor Odyssey CLx system.

To quantitatively determine cellular proteins (i.e. Supplemental Figure 1c), lysates were analyzed using the Protein Simple: an automated capillary-based protein separation and immuno-blotting assay. Lysates were prepared as above and diluted to 0.1 mg/mL with sample buffer and 40 mM DTT prior to loading 3 μL of sample onto each plate.

### Foci formation

Cells stably expressing doxycycline inducible short hairpins against XPR1 or the corresponding seed controls were plated at three different densities (18000, 12000, and 6000 cells/well) to determine optimal seeding density. Six replicates of each cell line were plated in 24 well plates, and half the wells were treated with 0.5 μg/mL doxycycline. Media and doxycycline were refreshed every two to three days until untreated wells reached 100 percent confluence. The cells were then washed with 1X PBS and fixed with 4% PFA in 1X PBS for 15 minutes. Fixation was quenched with deionized water and cells were stained with 0.1% crystal violet in deionized water for 20 minutes. Cells were then washed with deionized water to remove residual crystal violet and left to dry overnight. To quantify crystal violet staining, 10% acetic acid was incubated in each well for 20 minutes, diluted 1:3 with deionized water, and re-plated in quadruplicate in a 96 well plate. Absorbance was measured at 590 nm.

### In vivo sgRNA competition assay

A detailed protocol for the tumor formation competition assay is available online (https://www.protocols.io/view/in-vivo-nanopool-pooled-sgrna-competition-assays-t-bfsbjnan). A small lentiviral library of non-targeting, negative control (targeting gene deserts or introns), positive control (targeting pan-essential genes), and experimental sgRNA was made in an arrayed format and then pooled together. To minimize time in tissue culture, we optimized the infection and puromycin conditions to achieve roughly 30-50% infection efficiency and nearly 100% selection 50 hours after infection. OVISE cells were selected with 8 ug/mL Puromycin, and SNGM cells were selected with 4 ug/mL puromycin.

50 hours after infection, the cells were lifted, counted, and diluted in 50% matrigel at a final concentration of 8 million cells per 100 uL. Some cells were frozen to determine the early representation of the library. For in vitro experiments, the cells were re-plated and grown in standard conditions for 2 weeks. For in vivo experiments, mice were anesthetized under isoflurane gas, and two bilateral subcutaneous xenografts were inoculated in each of five mice (10 tumors per experiment). Tumors were measured twice weekly with calipers and tumor volumes were calculated using the formula: pi/6 x (width^2^ x length). Animal body weights were recorded weekly or twice weekly during the course of all studies. Mice were euthanized and tumors harvested on days 14, 21, and 28 after inoculation. After harvesting and weighing, tumors were flash frozen in liquid nitrogen until genomic DNA isolation.

At the end of the study, (28 days post inoculation; 30 days post-infection), the tumors were thawed, minced, and genomic DNA was extracted from all samples using the Qiagen DNeasy Blood and Tissue kits. Sample barcodes were sequenced by Illumina Next-Generation sequencing then deconvoluted with Broad Genetic Perturbation Platform’s PoolQ software for sgRNA read counts.

### Statistical Analysis for in vivo competition assay

The gene-level effect was determined by comparing normalized sgRNA read counts of the early time point (two days after infection, the day of inoculation) compared to the indicated time-points. If normalized read counts at the early time points were significantly different for a particular sgRNA, that sgRNA was not included in downstream analyses (as a sign that the plasmid did not produce lentivirus). sgRNA read counts were normalized such that the 7 “ cutting-control” sgRNA (representing viability effects from CRISPR/Cas9 genome editing) had a median depletion of 0. Next, the median fold change of all sgRNA targeting a particular gene was calculated, and only genes with 2-fold depletion or greater relative to the control sgRNA are reported per replicate in Supplemental Figure 2d-e. The different time-point replicates were median-averaged and are reported in Figure 1d. Statistical significance of XPR1 depletion was calculated by comparing the gene-level depletion of XPR1 (3 different sgRNA per replicate) compared to cutting control sgRNA (7 different sgRNA per replicate) using a t-test to compare all replicates at each time-point. The test was conducted in GraphPad Prism 8.0 and Holm-Sidak multiple comparisons correction was applied. Corrected p-values (q-values) are reported in Figure 1d. Test statistics are as follows: OVISE TC 2 weeks (n = 2, t-value = 29.6, 14 degrees of freedom); OVISE tumor 2 weeks (n = 4, t-value = 8.9, 30 degrees of freedom); OVISE tumor 3 weeks (n = 4, t-value = 7.1, 30 degrees of freedom); OVISE tumor 4 weeks (n = 2, t-value = 4.6, 30 degrees of freedom); SNGM TC 2 weeks (n = 2, t-value = 19.1, 14 degrees of freedom); SNGM tumor 2 weeks (n = 2, t-value = 5.6, 16 degrees of freedom); SNGM tumor 3 weeks (n = 4, t-value = 2.3, 28 degrees of freedom); SNGM tumor 4 weeks (n = 4, t-value = 5.0, 22 degrees of freedom).

### In vivo shXPR1_2 experiments in IGROV1

shRNA animal studies were approved by the Institutional Animal Care and Use Committee (IACUC) of the Broad Institute under animal protocol 0194-01-18. IACUC guidelines on the ethical use and care of animals were followed. IGROV1 cells with stable lentiviral integration of the indicated shRNA were secondarily infected with lentivirus encoding a eGFP-luciferase bicistronic construct as has been reported before 56. GFP positive cells were sorted using FACS. These cells were then inoculated into the intraperitoneal cavity of 7-week-old Rag1- /- Il2rg-/- (NRG) mice obtained from The Jackson Laboratories. Because of the pilot nature of this experiment, we tested four different cell densities (2, 4, 6, and 8 million cells) in duplicate for both the shXPR1_2 and shSeed_2 IGROV1 cell lines (16 total mice). Metastatic dissemination was monitored live as well as in individual organs ex vivo by bioluminescence imaging using the IVIS SpectrumCT (Perkin Elmer, Hopkinton, MA). For bioluminescence imaging, animals were injected with 150 mg luciferin/kg of body weight intraperitoneally. A luciferin kinetic study was performed for each tumor model to determine optimal imaging times for both live and ex vivo imaging. The animals were imaged live under isoflurane anesthesia. Photon and radiance emission was quantified using Living Image 4.5.1 Software.

Three weeks after significant tumor growth, we sorted the animals to control (Teklad Global 18% Protein Rodent Diet) or doxycycline (Teklad Global 18% Protein Rodent Diet containing 625 mg kg−1 doxycycline hyclate). At the time of treatment, one animal per inoculation group was treated with control or doxycycline chow. The two groups (-Dox/+Dox) displayed no difference in tumor burden at the time at the time of treatment (ANOVA with 3 degrees of freedom, F = 1.497, p = 0.26). Mice remained on their respective diets throughout the remainder of the study. Animal body weights were recorded twice weekly during the course of the study for body condition scoring. Animals were euthanized upon the development of ascites which occurred 2-3 weeks after treatment began.

We used a modified repeated measures ANOVA test to analyze the difference, if any, between the +Doxycycline and -Doxcycline groups in the IGROV1 shRNA in vivo experiment. We analyzed data up to day 15 post-treatment in which there were at least two animals in every group that had not reached a humane endpoint. Repeated measures ANOVA cannot handle missing values. We analyzed the data instead by fitting a mixed model as implemented in GraphPad Prism 8.0. This mixed model uses a compound symmetry covariance matrix, and is fit using Restricted Maximum Likelihood (REML). In the presence of missing values (missing completely at random), the results can be interpreted like repeated measures ANOVA. Fixed effects included in the model were time and treatment, with random effects of the individual mice. P-values evaluate the likelihood that, given repeated observations over time, the treatment group effects were observed by chance.

### Comparing expression of genes across normal and tumor tissues

We compiled log2(TPM + 1) gene expression data for normal fallopian tube (GTEx, n = 5), normal ovary (GTEx, n = 88), normal uterus (GTEx, n = 78), ovarian cancer (TCGA OV, n = 426) and uterine cancer (TCGA UCEC, n = 238) from the TOIL RSEM log2(TPM + 0.01) data at Xena Browser (https://xenabrowser.net/) and then converted the data to log2(TPM + 1). RNAseq gene expression for ovarian cancer cell lines (CCLE n = 40), and uterine cancer cell lines (CCLE, n = 22) were downloaded from the Cancer Cell Line Encyclopedia (https://depmap.org) as Log2(TPM + 1). Because most TCGA ovarian and uterine samples have relatively high purity (>80%)^57^, we used these data directly for the following comparisons. In Figure 2a and 2d and Supplemental Figures 4c and 4e, boxplots are drawn using the “ geom_boxplot” command in the R package ggplot2, such that the box spans the 1st and 3rd quartiles of values with the median indicated by a line. The whiskers extend 1.5x the interquartile range, and outlier’s beyond this range are excluded. In Figure 2a and 2d and Supplemental Figure 4c a Wilcoxon ranked sum test was employed using the R package ‘rstatix’ to compare the distribution of expression of the indicated gene between the indicated tissues, and p-values were corrected for multiple comparisons using Bonferoni’s method. In Supplemental Figure 4a a pairwise Wilcoxon ranked sum test was used to compare the expression of SLC34A2 in each tissue relative to all other tissues and p-values were corrected for multiple comparisons using Bonferoni’s method and are reported on the Y-axis. The difference in median SLC34A2 expression (tissue -expression across all tissues) is plotted on the X-axis of Supplemental Figure 4a. In Supplemental Figure 6c, the correlation between SLC34A2 and PAX8 mRNA was tested using a two-tailed pearson correlation test across the indicated tissues (n = 897). In Supplemental Figure 8c, the correlation between XPR1 and KIDINS220 mRNA was tested using a two-tailed pearson correlation test for the indicated tissue groups (all 60 tissues tissues, n = 17,194; top 15 correlated, n = 2,799, for all other tissues, listed from top to bottom, n = 337, 55, 172, 173, 36, 47, 182, 520, 66, 182, 496, 154, 119, 181, and 79).

### mRNA sequencing after PAX8 suppression

The indicated cell lines were obtained from CCLE and were modified to express dCas9-KRAB-BFP and the Tet Responsive element (TEG3G, Takara), the PAX8 overexpression constructs in vector pMT025^58^, and sgRNA from pXPR-BRD016. sgPAX8_4 (5’ -GGGGAACAAACTTCAGAAGG) suppresses PAX8 while sgPAX8_9 (5’ -GTGTGCAACCTCCGCCGTTG) does not. Three and six days after induction of dCas9-KRAB using doxycyline, RNA was extracted using a Qiagen RNEasy miniprep kit and then submitted to the Dana Farber Cancer Institute Molecular Biology Core Faciilty for library preparation, next generation sequencing, and analysis. mRNA levels were determined using cufflinks, and are presented as log2(FKPM + 1) The data presented in Figure 2d and Supplemental Figure 6e represent single samples collected at three days and are representative of the day 6 data (not shown). No statistical analysis was performed.

### Analysis of XPR1 copy number in TCGA

To evaluate the frequency of XPR1 amplification, we evaluated precomputed GISTIC2^28^ analyses for recurrent copy number alterations in ovarian and uterine TCGA datasets^17,18^. To compare the expression of XPR1 with its copy-number status, XPR1 copy number thresholds - as determined by GISTIC -were downloaded from CBIoPortal and then matched to the corresponding TCGA samples. 410 ovarian cancer and 171 uterine cancer samples were included in this analysis. In Figure 2b, each patient sample is represented by a horizontal line. Red indicates copy gain and blue indicates copy loss. Dashed vertical lines are the location of indicated genes. Data are a subset of the 489 samples with rank ordered by highest copy gain to indicate both focal and chromosome arm variants. In Figure 2c, we tested whether there was a significantly non-zero correlation between XPR1 copy number and XPR1 mRNA expression using a two-tailed spearman correlation test, and reported Spearman’s rho and the p-value. Also in figure 2c, the expression of XPR1 between tissue categories was compared using Wilcoxon ranked sums test with Bonferroni correction.

### Modifier Screen

The anchor modifier screen was performed as described previously^29^. OVISE cells stably expressing sgRNA targeting Chr2-2, XPR1_1, or XPR1-2 in the lentiviral guide-only vector pXPR-BRD016 were infected with the Brunello “ All-in-one” vector (pXPR-BRD023) in a format so that each cell received a maximum of one sgRNA. 24 hours post infection, the cells were split into two replicates and treated with 2 ug/mL puromycin to select for cells with stable integration. Every 3-4 days, the cells were trypsinized and re-plated to maintain a minimum representation of 500x library representation per replicate. After 15 days post-infection, all cells were collected and counted. The final representations for each replicate were as follows: 533x and 533x for sgChr2-2, 287x and 231x for the sgXPR1_1, and 81x and 83x for the sgXPR1_2. Genomic DNA was then extracted from each cell line and sgRNA barcodes were amplified by PCR.

### Modifier Screen Analysis

The representation of each sgRNA in each arm of the experiment was determined by next generation sequencing, and compared with plasmid DNA representation of the entire library. Using MAGeCK-MLE, this change in representation was converted to gene-level beta-values, representing the variability of each gene relative to all other genes. A permutation test (10 different permutations) on the sgRNA labels was used to empirically determine p-values, which were then corrected using the Benjamini-Hochberg Procedure^59^. Any gene with an FDR < 0.1 was considered a potential hit in that screen. SLC34A2 was the only statistically significant “ rescue” gene (i.e. a beta value > 0) for both sgXPR1_1 and sgXPR1_2 arms that was not significant in the control arm. There were several genes with beta-values significantly less than 0. Comparing the beta-values for this list of potential “ sensitizer” genes between the sgXPR1_1/2 arms and the sgChr2-2 arm, we note that all of these are significantly depleted across every arm, with little difference in beta-value. This indicates these are essential genes, and their depletion observed in every arm of the experiment is likely not due to a relationship with XPR1.

### Comparison of XPR1 dependency and tissue culture medium

Information on the growth medium is available for cell lines at depmap.org/portal. As each cell line is grown in a mixture of several medium types (e.g. 90% RPMI 1640 + 10% FBS), we estimated the concentration of phosphate by using a weighted average of phosphate concentrations of each component (see table below). The XPR1 dependency score was then compared to the concentration of phosphate using a one-tailed pearson correlation test in which we expected to find that more dependent cell lines were grown in higher concentrations of phosphate, and exact p-values and sample sizes are reported in Supplemental Figure 4a.

| Base | Manufacturer | Catalog # | [Phosphate] (mg/dl) |
| --- | --- | --- | --- |
| FBS | Various | Various | 5.3 |
| RPMI 1640 | Corning | 10-040-CV | 80 |
| DMEM | Thermo Fisher | 10566016 | 12.5 |
| Medium 199 | Thermo Fisher | 11150059 | 14 |
| McCoy's 5A | Thermo Fisher | 16600 | 58 |
| EMEM | ATCC | 30-2003 | 14 |
| MEM | Thermo Fisher | 51200038 | 14 |
| IMDM | Thermo Fisher | 12440079 | 12.5 |
| Ham's F-12 | Thermo Fisher | 21700075 | 14.2 |
| William's E Medium | Thermo Fisher | 12551032 | 14 |
| Waymouth's MB | Thermo Fisher | 11220035 | 64.2 |
| ACL4 | ATCC | MD-8002 | 0 |
| MEMa | Thermo Fisher | 32561037 | 15.8 |
| Leibovitz's L-15 | Thermo Fisher | 11415114 | 25 |

### Cell competition assays in low phosphate media

The protocol for luciferase-based cell growth competition assays was described in^58^. The basis of the experiment is to do a long term head to head competition assay between two variants of a cell line. Using lentivirus, each cell line was engineered to stably express one of two luciferase genes, either Firefly luciferase or Renilla luciferase. The Firefly luciferase expressing version of the cell line was also engineered to express Cas9.

For the competition assay in low phosphate, luciferase expressing cells were adapted from normal growth media (RPMI-1640 +10% Fetal Bovine Serum) to low phosphate media for 7 days. The concentration of phosphate in low phosphate media was empirically determined as the minimal phosphate concentration to support sustained cell viability for 3 weeks in culture. Low phosphate media was a 9:1 ratio of (RPMI-1640 without L-Glutamine and Phosphate +10% FBS; MP Biomedicals cat#: 09162975) with normal growth media (RPMI-1640 +10% Fetal Bovine Serum). We estimate 100% FBS to have 5.3 mg/dL of inorganic phosphate, though this value is likely to vary^60^. Therefore, this low-phosphate RPMI would have a concentration of 7.8 mg/ dL.

After cells were adapted to low phosphate media for 7 days, Renilla luciferase expressing cells were mixed at a 1:1 ratio with Firefly luciferase +Cas9 expressing cells and infected with the indicated sgRNAs on day 8. Cells were selected with puromycin 24 hours after infection. On day 11, cells were split and half were re-plated to propagate the cultured mixtures while the other half were subjected to Dual-Glo Luciferase Assay (Promega) to set baseline signal. The final luciferase assay was performed on day 16 after initial culture in low phosphate media.

### Measuring Intracellular Phosphate

To measure intracellular phosphate, cells were plated in 6-well plates as described above to perturb XPR1, KIDINS220, or SLC34A2. After the indicated times, the cells were washed three times with tris-buffered saline to remove residual phosphate from the media. The cells were then lysed in 1% NP-40, 50 mM Tris pH 7.5, and protease inhibitors, and the cellular debris was cleared by centrifugation. The cell lysate, or a dilution series of a phosphate standard, was then diluted in water to 50 uL in a 96-well microtiter plate and 10 uL Malachite Green Reagent A was added per manufacturer instructions (R&D Systems Cat# DY966). After 10 minutes, 10 uL of Malachite Green Reagent B was added to the samples and the absorbance at 620 nm was immediately read. If any sample was not within the linear range of the assay, then the samples were iteratively re-diluted and re-analyzed. Intracellular phosphate levels were calculated by dividing the interpolated phosphate concentration by the concentration of protein (determined by BCA assay) in each sample. In Figure 3e, Supplemental Figure 7a, and Supplemental Figure 9e, technical triplicates representative of at least 3 experiments are displayed.

### Immunofluorescence and fixed cell compatible dyes

Cells stably expressing V5-tagged XPR1 and KIDINS constructs were plated at a density of 10,000 -20,000 cells per well in NuncTM Lab-Tek™ II CC2™ 8-well Chamber Slides (Thermo Fisher). When investigating localization upon knockdown, cells were plated in chamber slides after 5 days of lentiviral transduction of sgRNA targeting XPR1, KIDINS220, or a non-coding portion of Chromosome 2. The following day, cells were washed with 1X Phosphate Buffered Saline (PBS) (Corning) and fixed in 4% paraformaldehyde (PFA) (Electron Microscopy Sciences) in 1X PBS for 15 minutes. Cells were washed twice with 1X PBS to quench fixation, permeabilized with 0.1% Triton X-100 in 1X PBS for 15 minutes, and were blocked in 1% bovine serum albumin (BSA) in 1X PBS for 1 hour. Cells were probed overnight at 4 °C with primary antibody diluted in 0.1% BSA in 1X PBS according to the table below. The following day, cells were washed three times with 0.1% Triton in 1X PBS and stained for 1 hour at 25 °C with Alexa-fluor conjugated secondary antibodies (Molecular Probes, ThermoFisher) diluted in the blocking buffer according to the table below. The wells were then washed with 1X PBS three times and counterstained with 4′,6-diamidino-2-phenylindole (DAPI) in 1X PBS at 2 μg/mL for 20 minutes. The wells were then washed twice with deionized water, and cells were mounted in ProLong Gold AntiFade Mountant (ThermoFisher).

For determining the organelle source of “vacuole-like” structures, the same general immunofluorescent staining protocol was used as above with the following changes. SNGM and OVISE Cas9 stable cell lines were plated at 10,000-20,000 cells per well on µ-Slide 8-well coated chamber slides (IBIDI, cat# 80826) and simultaneously infected with lentivirus expressing sgRNAs. The next day transduced cells were selected with 2 ug/mL puromycin for 24 hours, removed from puro selection and were fixed 6 days after infection. For ER labelling, cells were transduced with 24 uL of CellLight™ ER-GFP, BacMam 2.0 in 200 uL culture media 24 hours before imaging (Thermo-Fisher, cat# C10590) and no permeabilization step was performed. For mitochondrial imaging, cells were treated with 100 nM MitoTracker Red CMXRos (Thermo-Fisher, cat# M7512) in serum-free RPMI for 30 min at 37C, then MitoTracker dye media was replaced with normal growth media (RPMI with 10% FBS) and incubated for 1 hour at 37C before fixation. All other antibody-based stains were treated as described above. Information on the antibodies used and their concentrations for staining are provided in the antibodies Methods section.

### Multiplexed Transcriptional profiling

Multiplexed Transcriptional profiling (MixSeq^61^) was performed using custom pools of ovarian and uterine cancer cell lines. Cancer cell lines were pooled together (5-7 cell lines per mini pool) based on doubling time and frozen. To initiate the experiment, the cells were thawed and plated in 12-well dishes. The next day, virus encoding mixtures of sgRNAs (sgLacZ/sgChr2-2 or sgXPR1_1/sgXPR1_2) under conditions in which each cell received both sgRNA to increase the penetrance of inactivation. Cells were treated with 2 ug/mL puromycin 24 hours after infection. Four days after infection, the cells were lifted with TrypLE, spun down, resuspended in cell-staining buffer (PBS + 2%BSA + 0.02%Tween) and counted.

Perturbations were multiplexed for 10x sequencing using Cell Hashing^62^. Equal numbers of each ‘mini-pool’ were then pooled together, blocked with FcX blocking buffer (BioLegend) for 10 minutes on ice and then incubated with hash-antibodies (TotalSeq from BioLegend) for 30 minutes on ice. The cells were then washed thrice with cell-staining buffer and resuspended in Cell Capture buffer (PBS + 0.04% BSA), filtered with a 40 µm filter, and diluted to ∼1,500 cells per µL. A detailed protocol can be found online (https://www.protocols.io/view/cell-hashing-zn9f5h6). Approximately 40,000 cells were then loaded onto a 10x Chromium controller using v3. Single Cell 3’ reagent chemistries. Library preparation and next generation sequencing were performed as before^61^.

Sequencing data was processed using 10x Cell Ranger software (v3, hg19 reference genome), run with the ‘Cumulus’ cloud-based analysis framework^63^. SNP-based cell line classification and quality control was performed according to the methods described in reference 61. In brief, for each cell line the allelic fractions across a predefined 100,000 SNP reference set was estimated from bulk RNA-seq data using Freebayes^64^. A logistic regression model was then used to estimate the likelihood of the observed SNP reads for an individual cell having come from each cell line given the allelic fractions across the SNP reference set. A similar model was used to detect doublets where allelic fractions were modeled as a mixture of the allelic fractions from two different cell lines. After SNP-based classification, low quality cells and doublets were removed according to a set of stringent filters. These include the proportion of UMIs from mitochondrial genes being between 0.25 and 0.01, the number of reads at reference SNP sites being greater than 50, and the likelihood that a cell matches a particular cell line being much higher than the likelihood that it matches a different cell line or doublet.

MixSeq single-cell data was analyzed using the Seurat R package (v3^65^. The data was first normalized and scaled using the NormalizeData and ScaleData functions (default parameters). To generate the UMAP embedding (Supplemental Figure 7f), the top 5,000 most variable genes were selected using the FindVariableFeatures function, principal components were computed using the RunPCA function, and the embedding was generated using the RunUMAP function with 20 principal components, 10 nearest neighbors, and a minimum distance of 0.3 (default parameters otherwise). Cell cycle phase classification was performed with the CellCycleScoring function, using the S-and G2M-phase marker gene sets reported previously^66^. ΔG0/ G1 (Supplemental Figure 7g) was calculated for each cell line as the fraction of cells in G0/G1 in the XPR1 sgRNA condition minus the fraction of cells in G0/G1 in the control condition.

MixSeq differential expression analysis was performed using the “ limma-voom” pipeline^67,68^. First, per cell normalization factors were calculated using the “ TMMwzp” method from the edgeR R package^69^. Then, counts were converted into logCPM and the mean-variance relationship estimated using the voom function from the limma R package^68^. To identify each cell lines’ transcriptional response to XPR1 knockout removing the effect of cell cycle (Supplemental Figure 7g) a linear model was used with the ‘S.Score’ and ‘G2M.Score’ values from Seurat CellCycleScoring as covariates to regress out the effect of cell cycle. The top 500 differentially expressed genes displayed in the heatmap were identified based on the average log-fold-change across the 8 cell lines. To identify the average transcriptional response within the three highly correlated cell lines (RMGI, IGROV1, and OVISE) and five less correlated (JHUEM1,OVCAR4,COV413A, JHOS4, and HEC6) cell lines (Supplemental Figure 7f,g) a similar linear model was used, but cell line identify was added as a covariate to account for differences in baseline gene expression between the cell lines. P-values were derived from empirical-Bayes moderated t-statistics, and FDR adjusted Q-values were obtained using the Benjamini-Hochberg method.

### Phosphate uptake and efflux assays

To determine phosphate uptake or efflux, OVISE cells were infected in 6-well plates with lentivrius encoding sgRNA targeting XPR1, KIDINS220. Stable cell lines with SLC34A2 knockout were also used. The day before the experiment, the cells were split into 96-well plates such that the cells would be confluent the following day. Cells were first “ pulsed” using low-phosphate RPMI 1640 (a 1:9 ratio of standard RPMI to no-phosphate RPMI, see the lower phosphate competition assay above) supplemented with 10 μci/mL ^32^PO4 (Perkin Elmer NEX053001MC) and incubated at room temperature for 30 minutes. The cells were then washed with “ no phosphate” RPMI 1640. To determine phosphate uptake, and initial phosphate levels for time course efflux experiments, cells were lysed with 1% Triton X-100 and the amount of intracellular ^32^P was measured using a liquid scintillation counter. For efflux time course experiments, the “ chase’’ of phosphate efflux was then measured using high-phosphate RPMI (i.e. standard RPMI 1640). When incubated without phosphate in the medium (0 mM phosphate RPMI 1640), phosphate efflux is far lower (as has been reported before^9^ and this was used as a control. At each timepoint, the conditioned medium was taken, the cells were washed thrice with no-phosphate RPMI 1640 (to remove any radioactivity outside of the cells) and then lysed with 1% Triton X-100. Conditioned medium and cell lysates were analyzed for ^32^P using liquid scintillation counter. The extent of phosphate efflux was determined by dividing the ^32^P measured in the conditioned medium with the total 32P measured for that sample (in cell lysates and in the conditioned medium).

### High-throughput Mass Spectrometry analysis

High throughput protein interaction databases BioPlex^70^ and BioGRID^71^ were used to search and download lists of primary physical interactors of both XPR1 and KIDINS220. Interactions were identified by affinity capture-mass spectrometry, where epitope tags on target ‘bait’ proteins act as affinity capture probes for identifying ‘prey’ interactor proteins. Common interactors were found by comparing the four gene lists then visualizing the overlap.

### V5 CoIPs

HEK293T cells were transiently transfected with 40 μg of pLXTRC313 ORF vectors containing XPR1 constructs, KIDINS220 or GFP for 24 hours. Cells were then washed twice with 1X TBS and lysed with 0.4% NP40, 50 mM Tris, 150 mM NaCl, supplemented with Halt Protease Inhibitor Cocktail (ThermoScientific). Cell extracts were cleared by spinning at 15000 rpm for 10 minutes at 4 °C on a table top centrifuge, and quantified by BCA. 50 uL of V5-tagged immuno magnetic beads (MBL International, catalog number M167-11) was incubated with one milligram of protein lysate overnight at 4°C. The next day, beads were washed five times with 0.2% NP40 50mM Tris 150mM NaCl. Bound protein was then eluted by adding 10X sample buffer (Protein Simple) and incubating the beads for 30 minutes at 37°C. Eluate was removed by beads and was probed alongside whole cell lysates by Protein Simple Automated Western.

### Live cell lysotracker staining of acidic organelles

Similar to the immunofluorescence methods above, OVISE Cas9 stable cells were plated at 10,000 cells per well in µ-Slide 8 Well coated chamber slides (IBIDI, cat# 80826) and simultaneously infected with lentivirus expressing sgRNAs. 24 hours after infection, cells were selected with 2ug/mL puromycin for 2 days and then replaced with RPMI-1640 without phenol red supplemented with 10% FBS. Live cells were incubated with dyes and imaged 5 days after infection.

For dye staining, LysoTracker Red DND-99 (Invitrogen Cat#: L7528) was resuspended as a 1 mM stock in DMSO. Cells were stained in phenol-red free RPMI growth media 50 nM Lysotracker for 45 min at 37C in the dark. The dye was then washed out once with dye free growth media and then incubated with 1 ug/mL Hoechst 33342, (Invitrogen Cat#: H3570) in phenol red free growth media at 37C for 30 min and remained in Hoechst containing media during imaging. Cells were imaged immediately after on a Nikon Eclipse Ti microscope with a Yokogawa Life Sciences CSU-W1 spinning disc confocal system.

### Ultrastructural analysis by transmission electron microscopy

After infection with cutting control (sgChr2-2) or experimental sgRNA (sgXPR1_1 and sgKIDINS220_1), OVISE. Cas9 and EMTOKA cell lines were grown for 5 days in 6-well dishes. The cells were washed once with PBS and then fixed in 2.5% Glutaraldehyde and 2.5% Paraformaldehyde in 100 mM sodium cacodylate buffer, pH 7.4. Cells were then washed in 100 mM Sodium cacodylate buffer pH 7.4, postfixed for 30 min in a solution of 1% Osmium Tetroxide and 1.5% Potassium Ferrocyanide, washed in water thrice and incubated in 1% aqueous uranyl acetate for 30 minutes followed by two washes in water and subsequent dehydration in grades of alcohol (5 minutes each; 50%, 70%, 95%, twice in 100%).

Cells were removed from the dish in propylene oxide, pelleted at 3000 rpm for 3 minutes and infiltrated for 2 -16 hours in a 1:1 mixture of propylene oxide and TAAB Epon (TAAB Laboratories Equipment Ltd, https://taab.co.uk). The samples were subsequently embedded in TAAB Epon and polymerized at 60 °C for 48 hours. Ultrathin sections (about 60 nm) were cut on a Reichert Ultracut-S microtome, picked up on to copper grids stained with lead citrate and examined in a JEOL 1200EX Transmission electron microscope and images were recorded with an AMT 2k CCD camera.

## Supplemental Figures

**Supplemental Figure 1:**
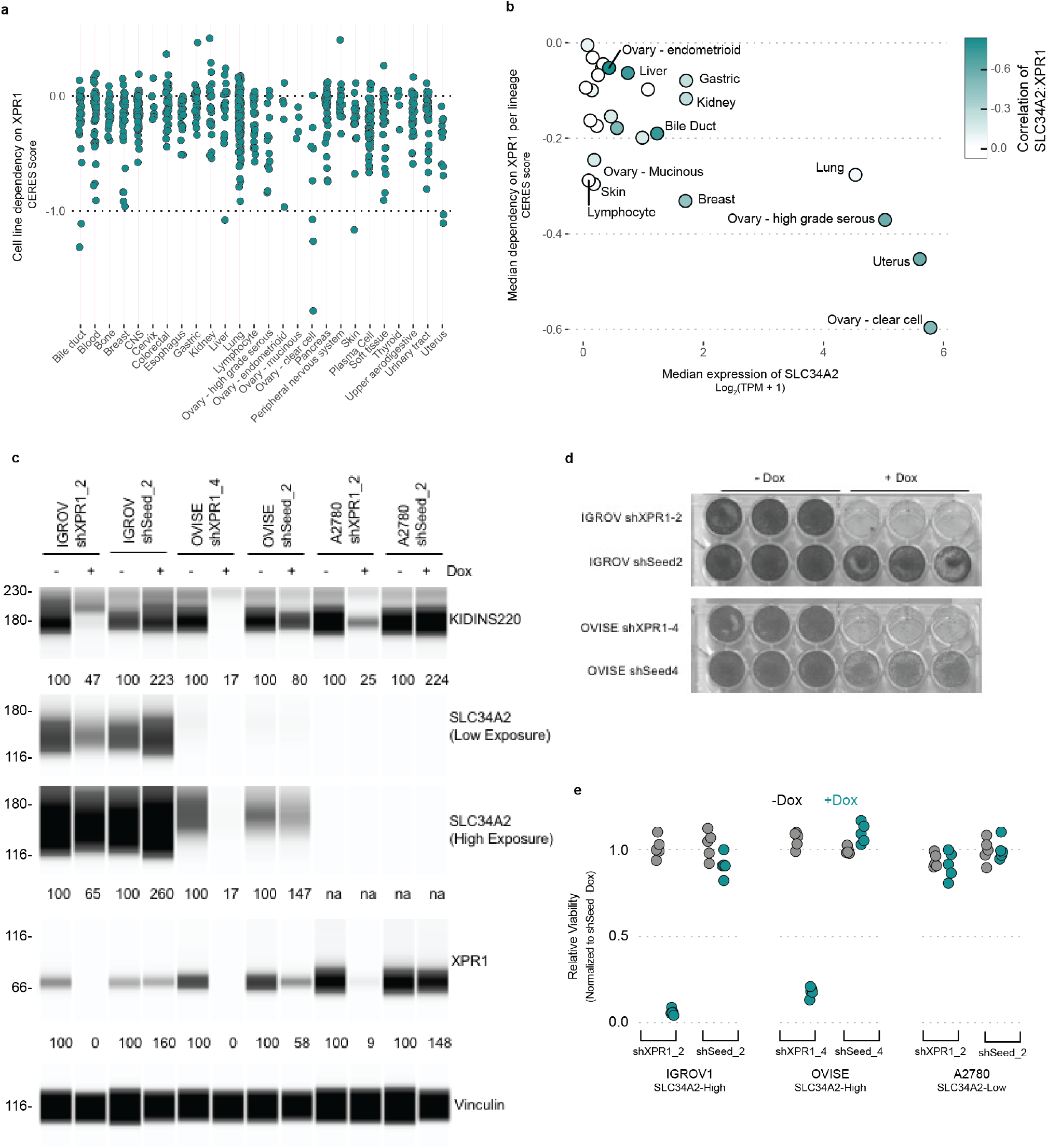
Validation of the XPR1 dependency in SLC34A2-overexpressing ovarian cancer cell lines using shRNA. **a**. For every cell line profiled in the Cancer Dependency Map dataset (739 cancer cell lines), the degree of XPR1 essentiality is plotted on the Y-axis. The CERES score is a scaled value of gene essentiality, where 0 is the effect of CRISPR/Cas9 genome editing and -1 is the effect of inactivation of pan-essential genes. Note that the ovarian lineage is separated into cancer subtypes. **b**. For every tissue type, the 10 highest SLC34A2 expressing cell lines were analyzed for their median expression of SLC34A2 (X-axis) and dependency on XPR1 (Y-axis). Note that some lineages may have less than 10 cell lines. Color encodes the correlation of SLC34A2 expression and XPR1 dependency across all cell lines within that lineage. **c**. Three days after induction of shRNA, protein levels were measured in cellular lysates by protein simple. Protein levels normalized to vinculin and the untreated (-Dox) conditions are expressed below each band. Note that shXPR1 reagents effectively suppress XPR1 protein levels but shSeed reagents do not. **d**. Colony formation assay to measure the long-term (14 day) penetrance and viability effect of suppression of XPR1 using shRNA in IGROV1 and OVISE cells. **e**. Seven day viability measured using Cell Titer Glo (Promega) and IGROV, OVISE and A2780 cell lines. IGROV1 and OVISE both express high levels of SLC34A2 and are predicted to be dependent on XPR1; A2780 does not.

**Supplemental Figure 2:**
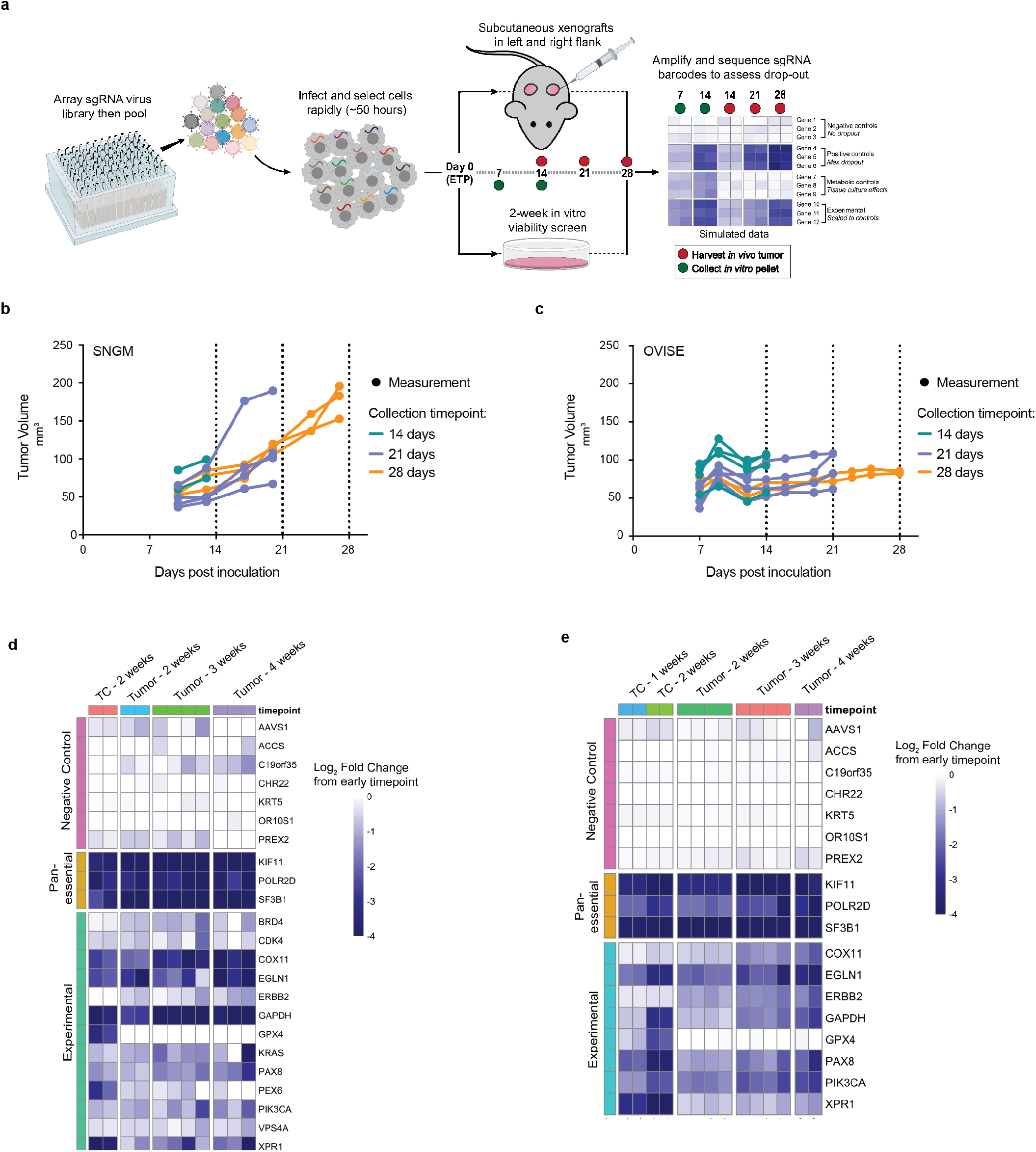
*In vivo* CRISPR/Cas9 competition assays for target validation in mouse xenografts. **a**. Experimental design for in vivo competition assays. Using a rapid infection and selection protocol, pooled sgRNA can be introduced via lentivirus into cancer cell lines and inoculated as subcutaneous xenograft and the effect of gene knock-out can be evaluated in a more physiologically-relevant environment than tissue culture. **b**. After rapid infection with pooled sgRNA, 8 million SNGM cells were inoculated as subcutaneous xenografts and allowed to grow. Tumor tissue was harvested at the indicated time points. **c**. Same as in b, but with the OVISE cancer cell line. **d**. sgRNA abundance in tumor xenografts was evaluated by PCR and next-generation sequencing analysis, and the fold change compared to the early time point is shown as a heatmap for all of the negative control genes as well as any gene with a >4 fold change in abundance in any of the screens. **e**. Same as in **d**, but with the OVISE cancer cell line.

**Supplemental Figure 3:**
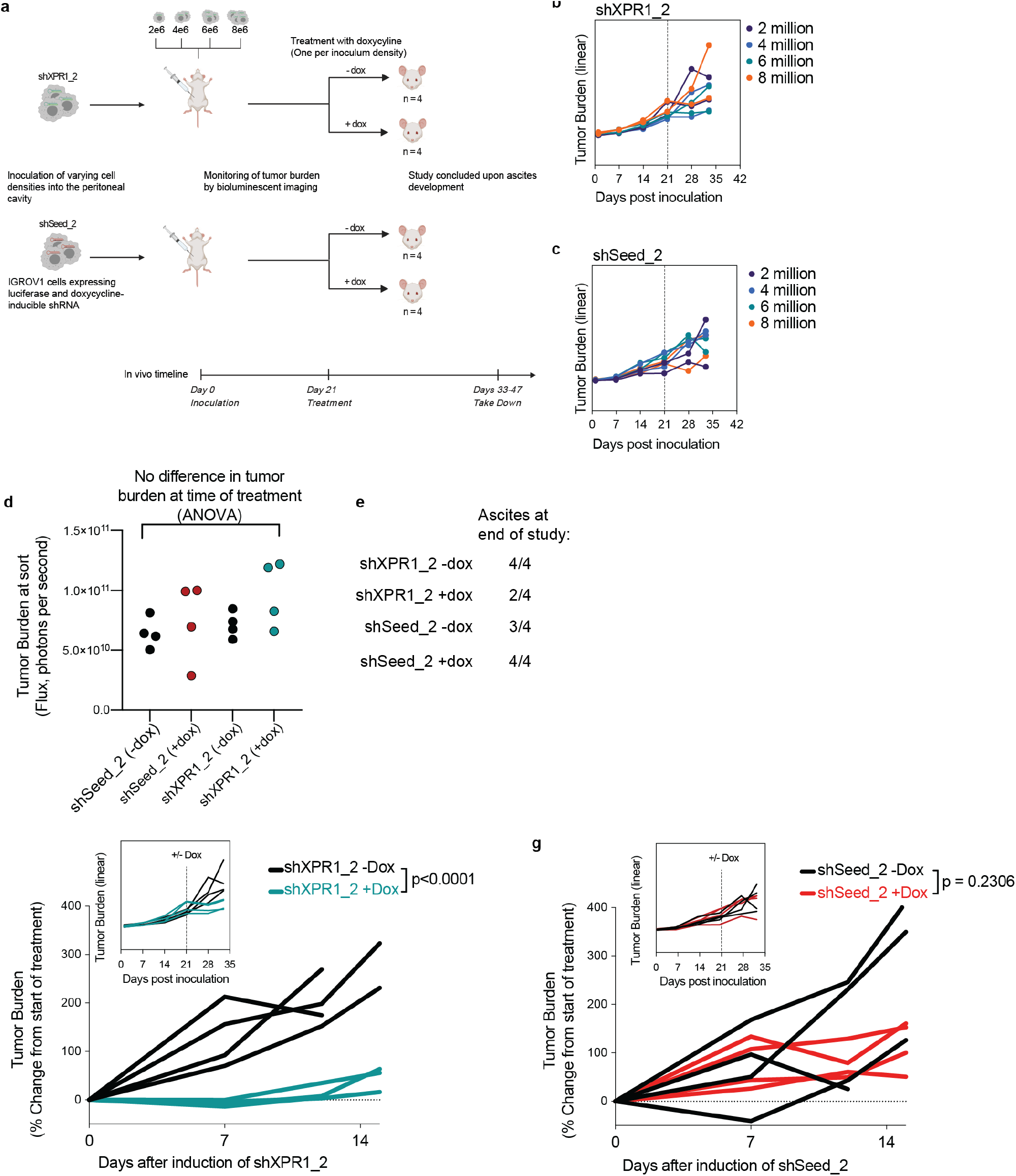
XPR1 suppression delays growth in disseminated ovarian carcinomatosis. **a**. Experimental design to assess the effect of XPR1 suppression on established intraperitoneal ovarian carcinomatosis. IGROV1 constitutively expressing luciferase were engineered to inducibly express shXPR1_2 (to suppress XPR1) or shSeed_2 (control). Because the in vivo growth kinetics for the model were unknown, 4 different cell densities were inoculated in the peritoneal cavity of mice, and tumor growth was monitored using bioluminescent imaging. After three weeks of tumor growth, one mouse from each inoculation group was fed with doxycycline chow to induce expression of shRNA. Animals developed ascites within 2-3 weeks after treatment, and so the study was terminated. **b**. The growth rate of IGROV1 cells engineered with shXPR1_2 was not dependent on the number of cells inoculated. **c**. Same as in panel **b**, but with IGROV1 cells engineered with shSeed_2. **d**. At the time of treatment, the tumor burden was equivalent across the four different groups. **e**. Number of animals per treatment and shRNA group which had developed ascites. **f**. The tumor growth of the SLC34A2 expressing ovarian cell line IGROV1, a model of disseminated ovarian carcinomatosis after XPR1 suppression using doxycycline-inducible shXPR1_2. The inset shows the full growth curves on a linear scale, while the larger image is the percent growth after treatment. **g**. Same as in **f**, but for the control shRNA (shSeed_2).

**Supplemental Figure 4:**
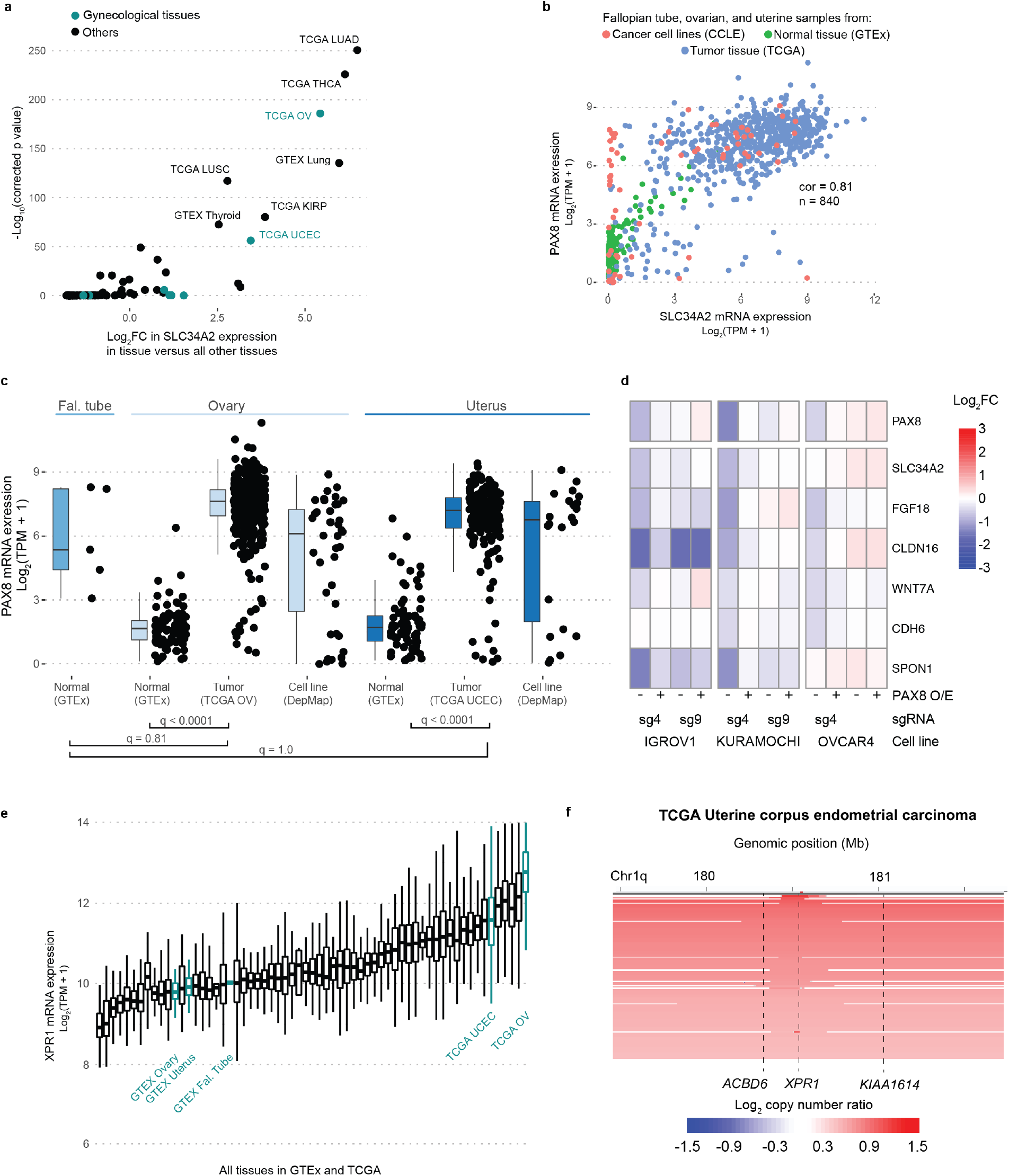
SLC34A2 and XPR1 overexpression in ovarian cancer are likely driven by PAX8**a**. Comparison of SLC34A2 expression across tissues. Using the combined GTEx, TCGA, and CCLE dataset, the differential expression of SLC34A2 in each tissue relative to the average of all tissues is compared. The relevant gynecological tissues (fallopian tube, ovary, and uterus) are highlighted in teal. The Cancer Genome Atlas abbreviations used include: LUAD = Lung adenocarcinoma; THCA = Thyroid carcinoma; KRP = Kidney renal cell papillary carcinoma ; LUSC = Lung squamous cell carcinoma; OV = Ovarian serous cystadenocarcinoma; UCEC = Uterine corpus endometrial carcinoma. **b**. The expression of PAX8 and SLC34A2 mRNA in the indicated tissues is plotted. The pearson correlation within these samples is indicated. **c**. PAX8 expression is compared from tissues in the GTEx, TCGA, and DepMap datasets as in Figure 2a and 2d. **d**. Gene expression -relative to parental cell lines profiled in parallel -of PAX8 target genes (see main text) after stable overexpression of WT-PAX8 (“ PAX8 O/E”) and/or induction of PAX8-target (sg4) or control (sg9) sgRNA and dCas9-KRAB. **e**. XPR1 expression across all tissues in TCGA and GTEx, with ovarian and uterine tissues highlighted in teal. **f**. XPR1 copy number heatmap for a ∼2.5 Mb region of chromosome 1 indicating XPR1 amplification in TCGA Uterine Corpus Endometrial Carcinoma 18. Each patient sample is represented by a horizontal line. Data are a subset of the samples rank-ordered by highest copy gain to indicate both focal and chromosome arm-level gains.

**Supplemental Figure 5:**
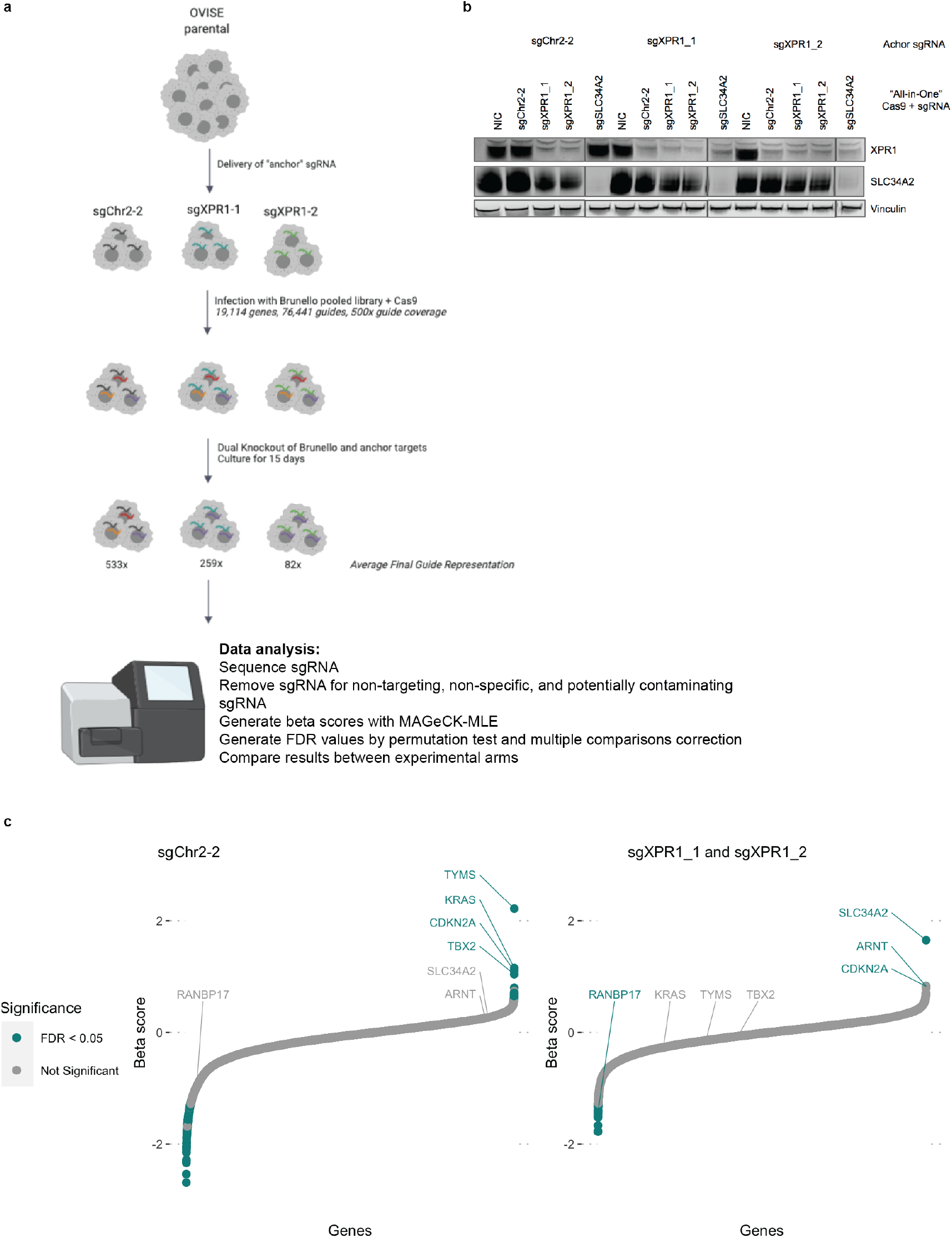
A genome-scale CRISPR/Cas9 screen validates the relationship between XPR1 dependency in the context of high expression of SLC34A2. **a**. Outline of the experimental method and analysis for a genome-scale, dual-knock-out modifier screen. OVISE (without Cas9 expression) is engineered to stably express sgRNA targeting XPR1. Upon introduction of “ all-in-one” lentivirus, containing both Cas9 ORF and a second sgRNA, both genes are simultaneously cut by Cas9. We used three sgRNA: one targeting a gene desert on chromosome 2 (sgChr2-2) and two targeting XPR1 (sgXPR1_1 and sgXPR1_2) and infecting the cells with the Brunello genome-scale sgRNA library. 15 days after infection, we collected cell pellets (and sequenced the sgRNAs. **b**. Western confirmation of dual-knock-out of XPR1 and SLC34A2. The three cell lines used in the genome-scale screen were infected with “ all-in-one” lentivirus expressing control-, XPR1-, or SLC34A2-targeting sgRNA. Note that in the sgXPR1 “ anchor” cell lines, XPR1 is suppressed with the control virus, indicating that the first infection provides XPR1-targeting sgRNA and the second infection provides Cas9 protein. NIC stands for “ no-infection control’. **c**. Arm-level results of the genome-scale modifier screen. See methods for full analysis details. Beta-scores represent the extent to which a gene was enriched or depleted relative to the initial plasmid representation. An XPR1-positive and control-neutral score represents a likely rescue gene (i.e. SLC34A2 and ARNT). XPR1-positive and control-positive scores represent genes with profound viability defects without specificity for XPR1 (e.g. RANBP17).

**Supplemental Figure 6:**
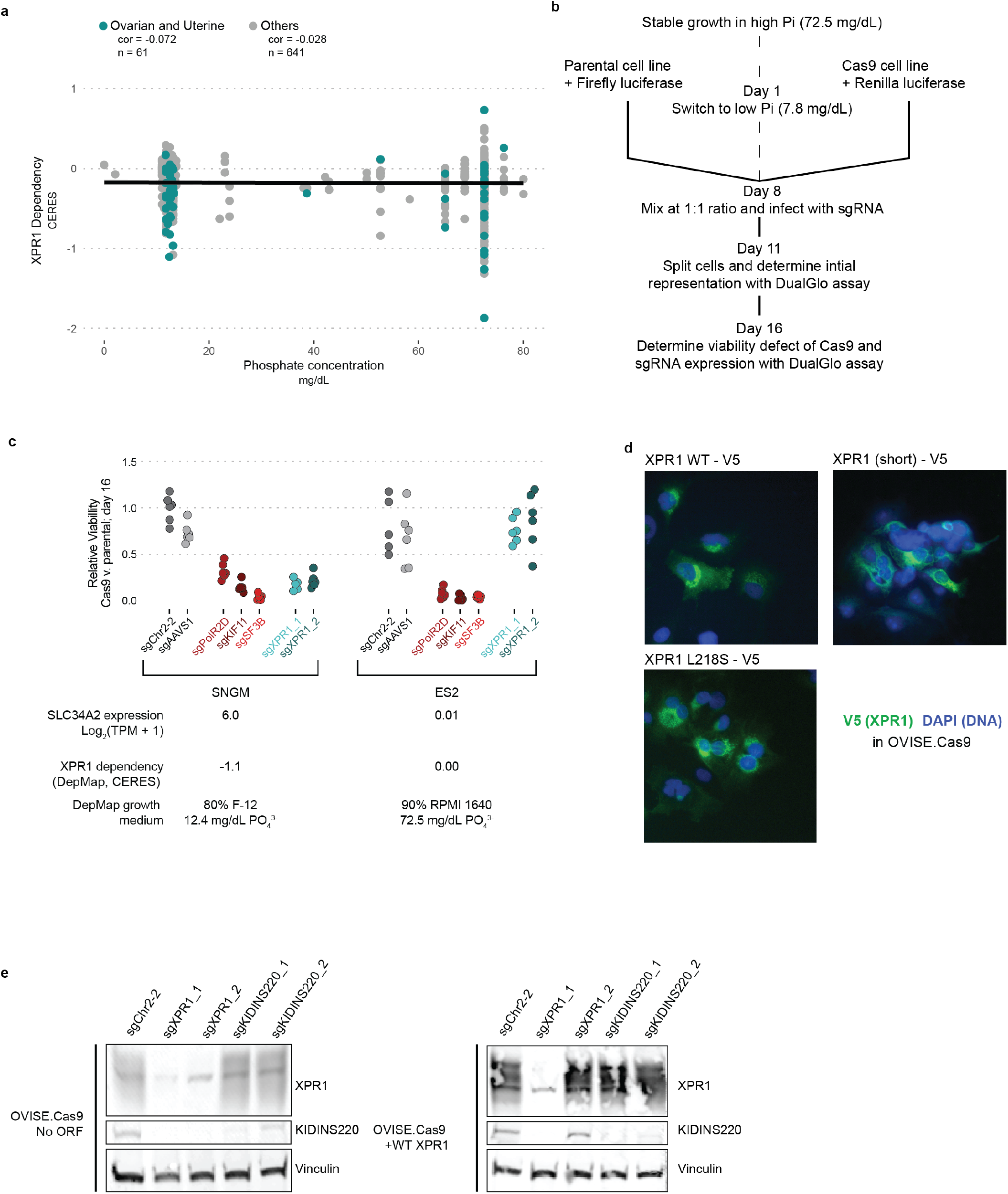
The XPR1 dependency is not affected by phosphate levels in the tissue culture medium. **a**. The concentration of phosphate in the growth medium of DepMap cell lines does not determine the extent of XPR1 dependency. Concentrations of phosphate were estimated from manufacturer formulations (see methods) and the pearson correlation between growth medium phosphate and XPR1 dependency is indicated. **b**. Experimental procedure for manipulating tissue culture medium and assessing its effect on XPR1 dependency. The same parental cancer cell line was engineered to express firefly luciferase and Cas9, or renilla luciferase alone. After a one-week adaptation to lowered phosphate, the two variants were mixed together and infected with sgRNA-encoding lentivirus. After selection for lentivirus-infected cell lines, the initial representation of Cas9:parental cells was determined by measuring the ratio of Firefly:Renilla luciferase using a DualGlo assay (Promega). One week after infection (Day 16 of the protocol), the extent to which genetic perturbation was detrimental to cell viability was determined using the DualGlo assay. **c**. The XPR1 dependency is maintained in a low phosphate medium condition. SNGM and ES2 were profiled in the assay outlined in panel b. Note that the CERES score -displayed below the plot -represents the viability defect of the cell line 21 days after knock-out of XPR1 and growth in the indicated growth medium. **d**. Immunofluorescent localization of XPR1 mutants using the V5 epitope tag. Left, WT XPR1 localizes to the secretory pathways as well as puncta within the cytoplasm. Middle, XPR1 (short) staining appears more diffuse. The L218S mutation has the same localization pattern as WT XPR1, consistent with proper trafficking, similar to what has been observed previously (see main text). **e**. Western blot validation of guide-resistant ORF. OVISE.Cas9 cells (parental, left, or overexpressing the WT XPR1 ORF, right, used in Figure 3e) were infected with the indicted sgRNA and harvested 5 days after infection. The XPR1 ORF includes a mutation to block binding of sgXPR1_2 but not sgXPR1_1. Note the inactivation of both endogenous and overexpression ORF with sgXPR1_1 and only endogenous XPR1 with sgXPR1_2.

**Supplemental Figure 7:**
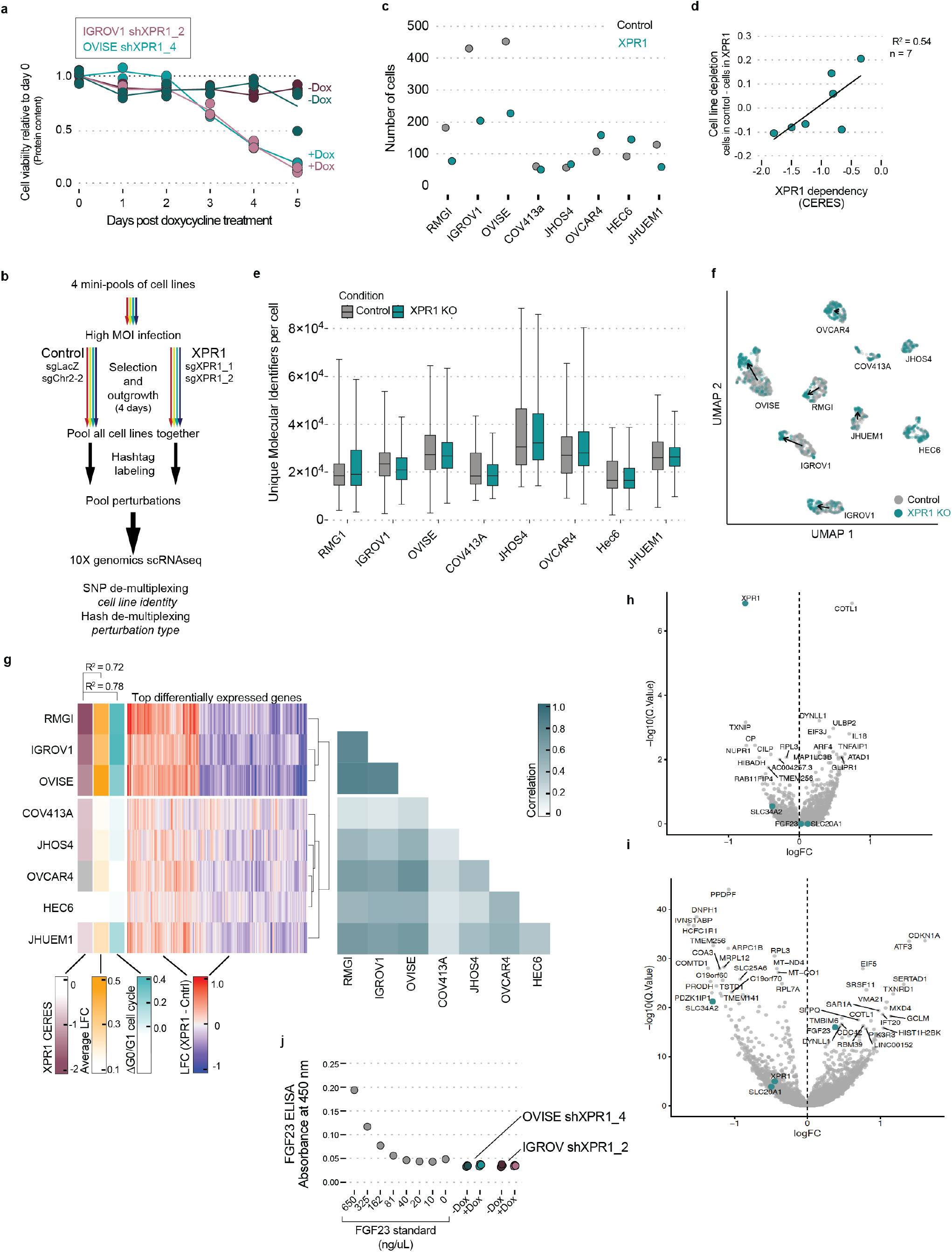
Transcriptional profiling reveals a phosphate-related homeostatic response after XPR1 inactivation. **a**. The viability of cells (as measured by total protein content) was measured in parallel with total phosphate as in Figure 3e. **b**. Schematic for the MixSeq determination of transcriptional effects after XPR1 inactivation. **c**. TThe total number of cells observed in the MixSeq experiment. **d**. The difference in number of cells observed in panel **c. e**. The total number of unique transcripts measured for each cell, as measured by unique molecular identifiers (UMIs). **f**. UMAP projection of the 2,501 cells from the indicated cell lines (determined by SNP profiles) and perturbations (indicated by cell-surface antibody ‘hash-tag’ labeling). **g**. Middle, the log-fold change of the top 500 differentially expressed genes after regressing out the effect of cell cycle. Left, summary annotations for each cell line include XPR1 dependency (XPR1 CERES), the overall transcriptional change (average log2 fold-change), and the degree of cell cycle arrest observed in the single-cell data (ΔG0/G1). Right, the pearson correlation of the top 500 differentially expressed genes between each cell line. **h**. Differentially expressed genes -after correcting for cell cycle -in the less sensitive cell lines (COV413a, JHOS4, OVCAR4, HEC6, and JHUEM1). **i**. Same as in **h**, but for the highly correlated cell lines RMG1, IGROV1, and OVISE. **j**. Four days after induction of shXPR1_2 (IGROV1) or shXPR1_4 (OVISE) using doxycycline, the amount of secreted FGF23 was measured in the conditioned medium using ELISA.

**Supplemental Figure 8:**
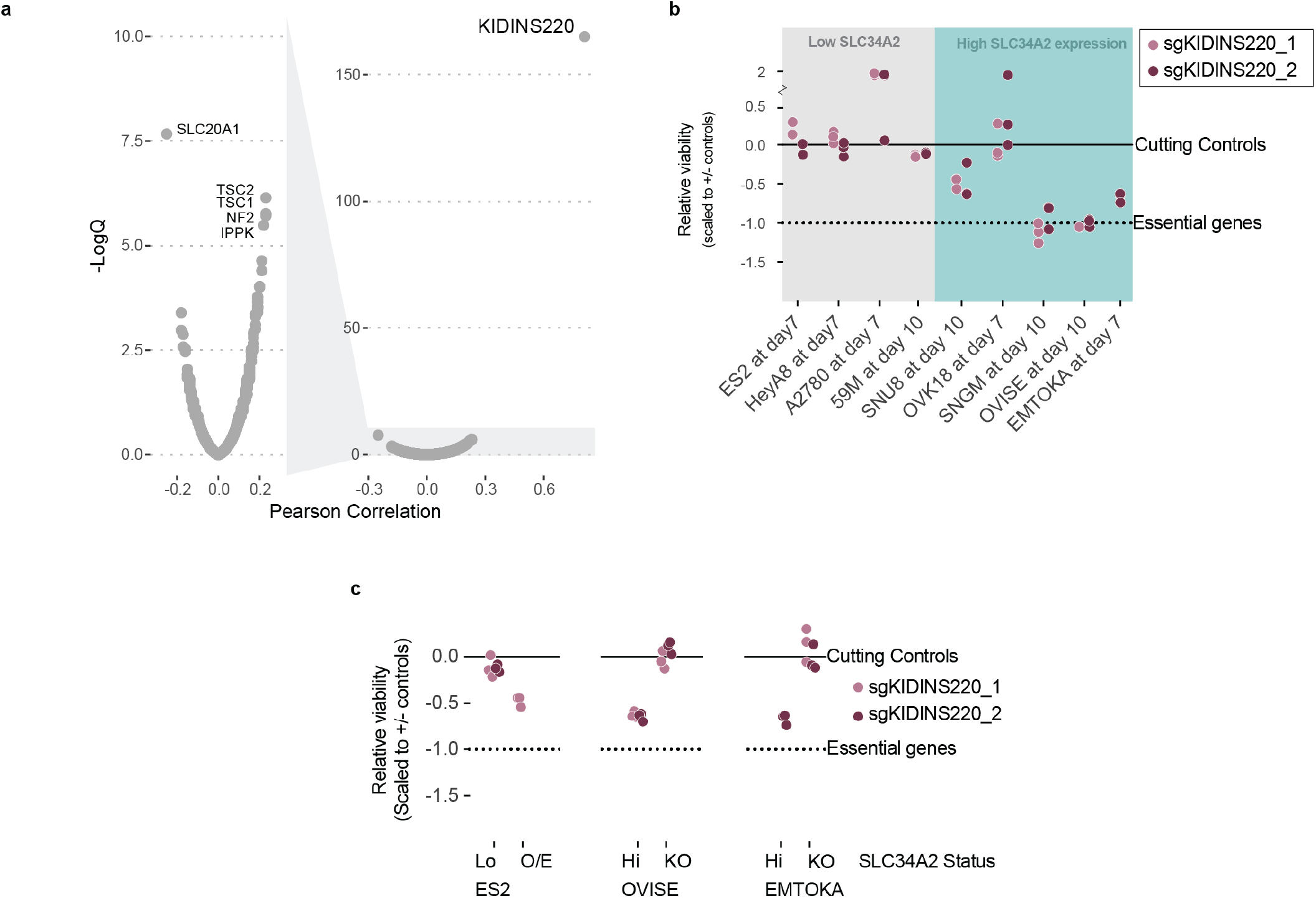
KIDINS220 dependency is highly correlated with XPR1 dependency. **a**. Volcano plot for the correlation of XPR1 dependency with all other genes assayed in large scale CRISPR/Cas9 loss of viability screens across 739 cancer cell lines. **b**. Validation of KIDINS220 dependency was performed in ovarian and uterine cancer cell lines with a range of SLC34A2 expression as in Figure 1c. **c**. SLC34A2 expression is necessary and sufficient for KIDINS220 dependency, performed as in Figure 3b.

**Supplemental Figure 9:**
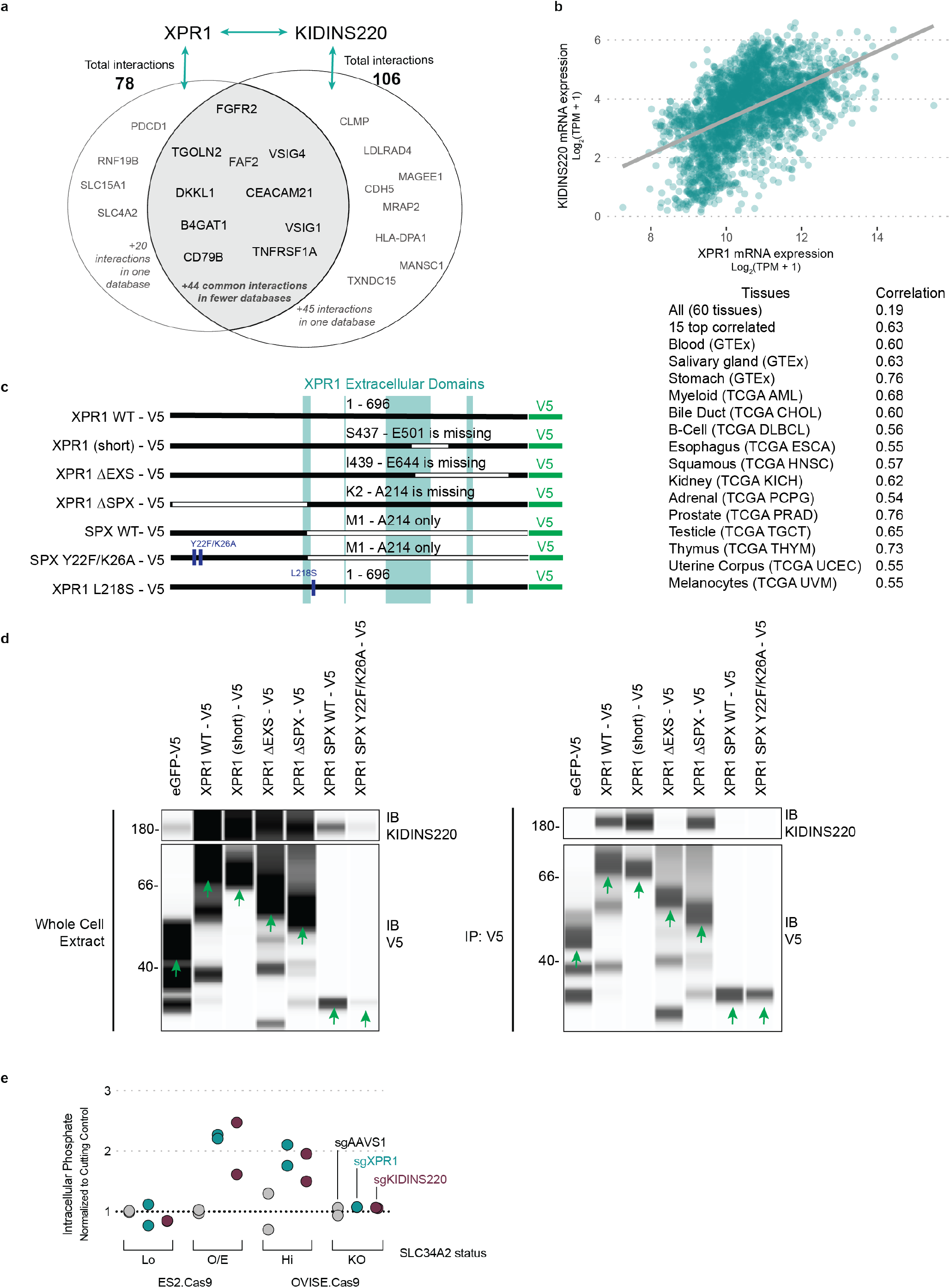
XPR1 and KIDINS220 are members of a co-regulated phosphate export complex. **a**. High throughput protein:protein interaction databases implicate XPR1 and KIDINS220 as part of a protein complex. The interacting partners of XPR1 and KIDINS220 were downloaded from the BioGrid and Bioplex databases and compared. Proteins present in the XPR1 or KIDINS220 interactomes are highlighted as text. **b**. Top, the mRNA expression of XPR1 and KIDINS220 is shown for the fifteen tissues with the highest correlation in expression. The line represents linear regression for these samples (n = 2,799). Bottom, the Pearson correlation for those tissues, highlighting the diverse tissues in which there is a high correlation between XPR1 and KIDINS220 expression. **c**. Mutants of XPR1 used in this manuscript. XPR1 WT refers to the 696 amino acid protein produced by NM_004736 (the only isoform detected by RT-PCR of OVISE mRNA), while XPR1 (short) refers to the 631 amino acid product of NM_001135669. All constructs have C-terminal V5 tags for immunoprecipitation, western blotting, and immunofluorescent detection. **d**. Right, co-immunoprecipitation of XPR1 domain mutants with endogenous KIDINS220 in 293s. This is an uncropped version of Figure 3b, including the SPX domain only constructs (lanes 6-7). Left, the whole-cell extract from the experiment, indicating differing expression levels of KIDINS220 or XPR1 constructs. It should be noted that much of the background signal is attributed to immunoblotting for endogenous XPR1 in the same experiment. Green arrows indicate the expected molecular weight of the ORF, in contrast to degradation products. **e**. Five days after infection with the indicated sgRNA targeting XPR1 or KIDINS220, free inorganic intracellular phosphate was determined as in Figure 3e.

**Supplemental Figure 10:**
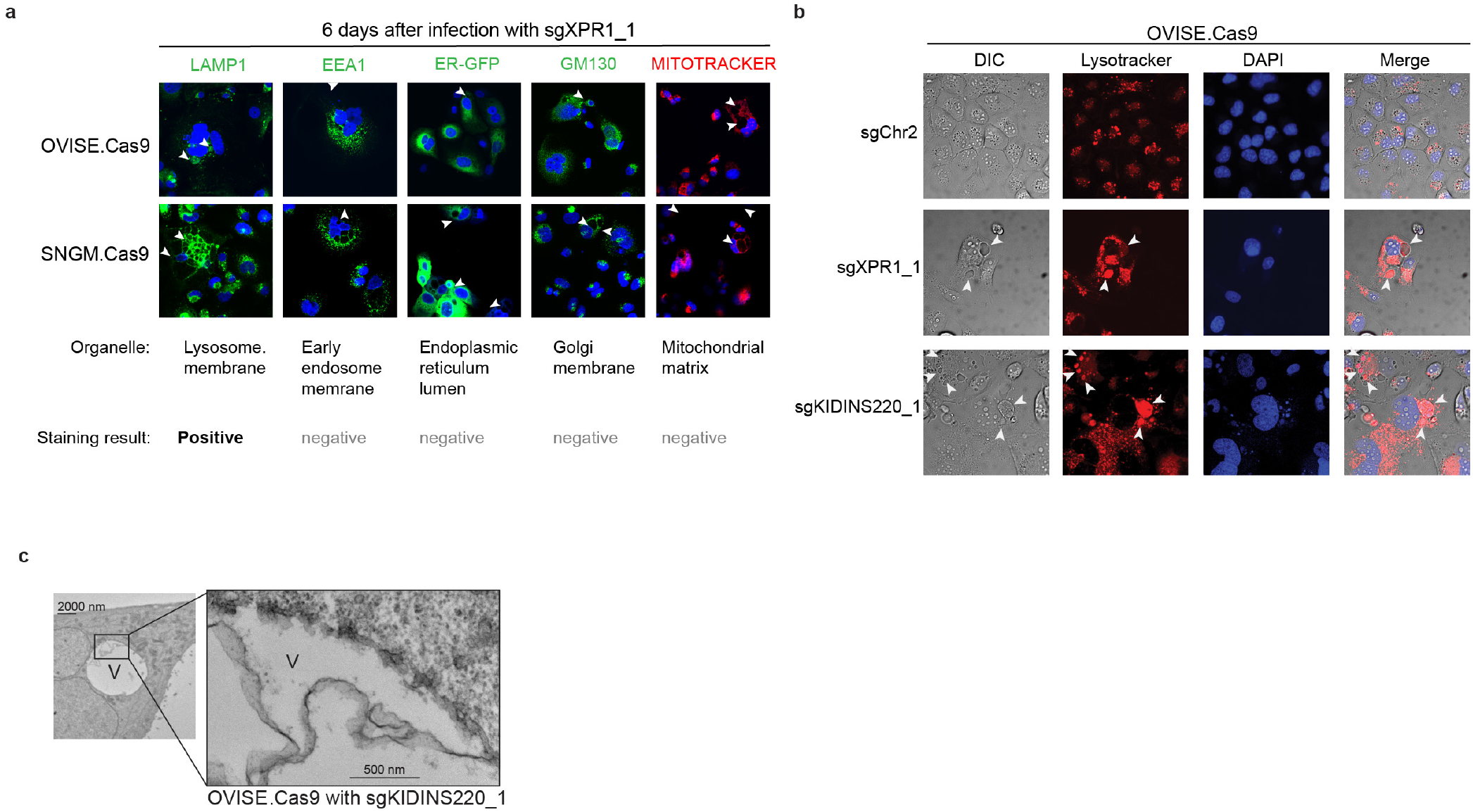
Vacuole structures precede cell death and are not derived from many common organelles. **a**. 66 days after infection with lentivirus encoding sgXPR1_2, OVISE.Cas9 and SNGM.Cas9 cell lines were stained and imaged using the indicated dyes and stains. Arrowheads indicate the location of vacuole structures by phase contrast (not pictured). Positive staining was only observed for the lysosomal dye LAMP1. **b**. Live cell DIC and confocal immunofluorescence images of XPR1 dependent cell line OVISE 5 days after CRISPR inactivation of XPR1 or KIDINS220 vs control sgRNA (sgChr2-2). Acidic organelles were detected with Lysotracker (red) and DNA with DAPI (blue). Arrowheads indicate the location of ‘vacuole-like’ structures. **c**. Ultrastructural analysis of OVISE cells 5 days after infection with sgKIDINS220_2, as in Figure 4g.

